# Procyanidin C1 is a natural agent with senolytic activity against aging and age-related diseases

**DOI:** 10.1101/2021.04.14.439765

**Authors:** Qixia Xu, Qiang Fu, Zi Li, Hanxin Liu, Ying Wang, Xu Lin, Ruikun He, Xuguang Zhang, Judith Campisi, James L. Kirkland, Yu Sun

## Abstract

Aging causes functional decline of multiple organs and increases the risk of age-related pathologies. In advanced lives, accumulation of senescent cells, which develop the senescence-associated secretory phenotype (SASP), promotes chronic inflammation and causes diverse conditions. Here we report the frontline outcome of screening a natural product library with human primary stromal cells as an experimental model. Multiple candidate compounds were assayed, and grape seed extract (GSE) was selected for further investigation due to its leading capacity in targeting senescent cells. We found procyanidin C1 (PCC1), a polyphenolic component, plays a critical role in mediating the antiaging effects of GSE. PCC1 blocks the SASP expression when used at low concentrations. Importantly, it selectively kills senescent cells upon application at higher concentrations, mainly by enhancing production of reactive oxygen species (ROS) and disturbing mitochondrial membrane potential, processes accompanied by upregulation of Bcl-2 family pro-apoptotic factors Puma and Noxa in senescent cells. PCC1 depletes senescent cells in treatment-damaged tumor microenvironment (TME) and enhances therapeutic efficacy when combined with chemotherapy in preclinical assays. Intermittent administration of PCC1 to both senescent cell-implanted mice and naturally aged animals alleviated physical dysfunction and prolonged post-treatment survival, thus providing substantial benefits in late life stage. Together, our study identifies PCC1 as a distinct natural senolytic agent, which may be exploited to delay aging and control age-related pathologies in future medicine.

## Introduction

Aging is considered the single largest risk factor for multiple chronic disorders such as metabolic disorders, cardiovascular diseases, neurodegenerative pathologies and diverse types of malignancies, which together account for the bulk of morbidity, mortality and health costs globally ^1^. Considerable progress is made over recent years in development of specific agents to treat individual aging-related conditions, including type 2 diabetes, osteoporosis, skeletal fragility and vascular dysfunction. However, the combined effect of these drugs on controlling morbidity and mortality of chronic diseases has been modest, which tend to occur in synchrony as multimorbidities, with a prevalence increasing exponentially after 70 years of age ^2^.

Several major factors affecting healthspan and lifespan have been identified through studies across a range of species, and defined as aging mechanisms that can be categorized into nine hallmarks ^3^. Of these fundamental aging mechanisms, cellular senescence has received substantial attention, as it represents a process typically druggable to prevent or delay multiple aging comorbidities ^4^. First reported in 1960s, cellular senescence is a cell fate involving essentially irreversible replicative arrest, profound chromatin changes, apoptosis resistance and increased protein synthesis, culminating in overproduction of pro-inflammatory cytokines, a feature termed the senescence-associated secretory phenotype (SASP) and responsible for aging and myriad age-related pathologies ^5^. Prophylactic ablation of senescent cells positive for the senescence marker p16^INK4A^ mitigates tissue degeneration and extends animal healthspan, supporting that senescent cells play a causative role in organismal aging ^6, 7^.

While much remains to be done, success in preclinical studies has inspired the initiation of proof-of-concept clinical trials involving senolytics for several human diseases and holding the potential to decrease the burden of *in vivo* senescent cells via selective elimination with pharmacological agents ^8–10^. Since the first discovery in 2015 ^11^, a handful of synthetic or small molecule senolytic agents are reported, with prominent capacity demonstrated in eradicating senescent cells. Targeting strategies were mainly based on the resistance mechanism of senescent cells to apoptosis, which emerge as senescence-associated anti-apoptotic pathways (SCAPs) and allow senescent cells survival for an extended period ^12, 13^. Of note, intermittent administration of senolytics holds the potential to reduce the risk of patients developing adverse conditions, minimize off-target effects of drugs and prevent development of drug resistance of senescent cells, which do not divide and differ from cancer cells, as the latter frequently acquire advantageous mutations against anticancer therapies. However, most of reported senolytics are cell lineage or cell type-dependent, or alternatively, exhibit substantial cytotoxicity *in vivo*, thus limiting their potential use for clinical purposes.

In this study, we screened a natural product medicinal library composed of anti-aging agents and identified several candidate compounds including grape seed extract (GSE). Further analysis revealed that PCC1, a B-type trimer epicatechin component of GSE flavonoids, plays a major role in inhibiting SASP expression at low concentrations and killing senescent cells at high concentrations, the latter case via apoptosis induction. Preclinical data suggested that upon combination with classic chemotherapy, PCC1 can significantly reduce tumor size and prolong experimental animal survival. Thus, our study consistently supports that PCC1 can be exploited as a safe geroprotective agent, and more specifically, a novel phytochemical senolytic isolated from natural sources, to delay aging and curtail age-related diseases, thus providing clues for exploration of its clinical merits in geriatric medicine.

## Results

### GSE restrains the SASP expression when used at low concentrations

In an effort to identify new compounds that can effectively modulate senescent cells, an unbiased agent screening was performed with a phytochemical library composed of 45 plant-derived medicinal agents (PDMA library). We chose to employ a primary normal human prostate stromal cell line, namely PSC27, as a cell-based model for this purpose. Composed mainly of fibroblasts but with a minor percentage of non-fibroblast cell lineages including endothelial cells and smooth muscle cells, PSC27 is primary in nature and develops a typical SASP after exposure to stressors such as genotoxic chemotherapy or ionizing radiation ^14–17^. We treated these cells with a pre-optimized sub-lethal dose of bleomycin (BLEO), and observed increased positivity for senescence-associatedβ-galactosidase (SA-β-Gal) staining, decreased BrdU incorporation, and elevated DNA damage repair (DDR) foci several days afterwards (Supplementary Fig. 1a-c). We set up a screen strategy to compare the effects individual medicinal products generated on the expression profile of senescent cells (Fig. 1a).

**Fig. 1.**
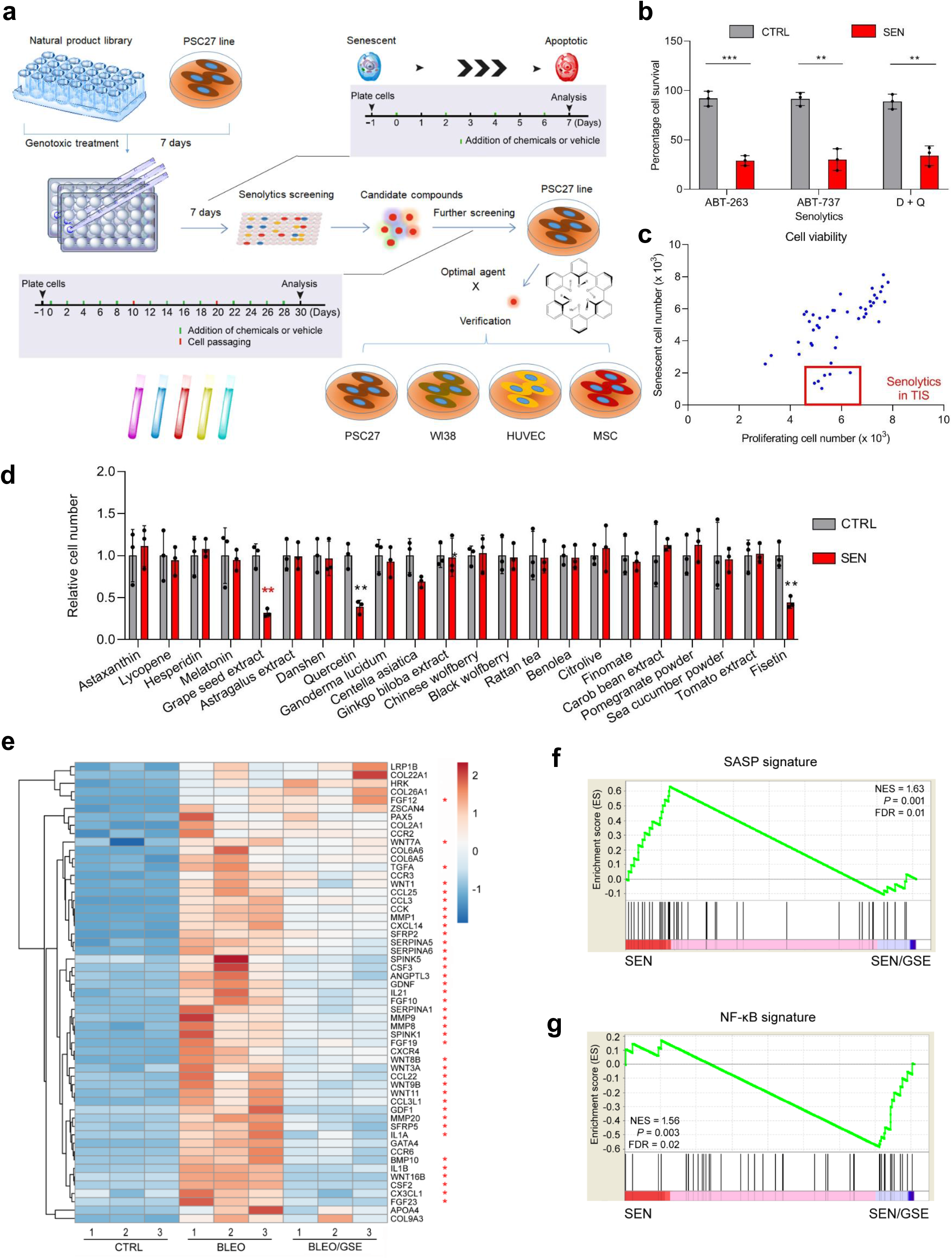
*In vitro* screening of a natural product library with human stromal cells. (**a**) Schematic workflow of cell-based screening strategy for a natural library of medicinal agents. (**b**) Quantification of cell survival of control (CTRL) and senescent (SEN) PSC27 after treatment with ABT-263 (1 μM), ABT-737 (10 μM) or Dasatinib (250 nM) combined with Quercetin (50 μM) for 3 d. (**c**) Screening results after pharmacological assays. Library agents were assessed at a concentration of 3 μg/ml with 5.0 × 10^3^ cells for 3 d. Hits were selected based on their ability to specifically kill senescent cells. Each blue dot represents a single agent as the mean of three replicates. Red rectangular region denotes potential candidates of senolytics in the case of TIS. (**d**) Appraisal of the effects of randomly selected agents (22 representative candidates out of 45 in the library) on the survival of CTRL and SEN cells. (**e**) Heatmap depicting the expression profiles of human genes in CTRL cells, SEN cells or SEN cells treated by GSE. According to the fold change of transcript expression, top 50 genes are displayed. (**f**) GSEA blot of significant gene set indicative of a typical SASP. (**g**) GSEA blot of significant gene set associated with the activation of NF-κB signaling. Cellular senescence was induced by the genotoxic chemical bleomycin (BLEO), which typically induces the TIS. **, *P* < 0.01; ***, *P* < 0.001. TIS, therapy-induced senescence. Data in **b-d** are shown as mean ± SD and representative of 3 independent biological replicates.

One promising advantage of senolytic agents is to selectively induce programmed death of senescent cells, such as ABT-263, ABT-737 and combined use of dasatinib and quercetin (or alternatively, ‘D + Q’) ^11, 18, 19^. We first tested the efficacy of these geroprotective drugs against senescent PSC27, to demonstrate its potential as an experimental cell model for drug screening. Our preliminary data suggest that each of these compounds was able to significantly deplete senescent, but not proliferating cells, thus confirming the feasibility of using this stromal line for further studies (Fig. 1b). Upon large scale screening of the PDMA library, we noticed that several compounds displayed the potential to selectively kill senescent cells in culture (Fig. 1c).

Among the agents showing preliminary anti-senescence effects, we first noticed GSE, quercetin and fisetin, as each displayed a prominent efficacy in senescent cell elimination (Fig. 1c). Since quercetin and fisetin share similar chemical structure, exert similar medicinal effects, and are both reported as senolytics ^11, 20, 21^, we chose to focus on GSE, which remained a largely underexplored source. Under *in vitro* condition, we treated PSC27 cells at a series of GSE concentrations. The data showed that at 0.1875 μg/ml GSE suppressed the SASP with a maximal efficiency. However, lower or higher concentrations of this agent achieved less efficacy, though the latter may be presumably linked with cell stress response induced by increased cytotoxicity (Fig. 1d). We then sought to profile genome-wide gene expression via unbiased assays. Data from RNA-seq indicated that treatment with GSE significantly altered the expression profile of senescence cells, with 2644 genes downregulated and 1472 genes upregulated with a fold change of 2.0 *per* gene (*P* < 0.01) (Extended Data Fig. 1a). Although expression of a few SASP-unrelated genes showed a similar tendency as those typical SASP factors (Fig. 1e), data from GSEA analysis consistently supported restrained expression of molecular signatures reflecting expression of the SASP or activation of the NF-κB complex, the latter a well-established event mediating development of the pro-inflammatory phenotype (Fig. 1f,g). Protein-protein interaction (PPI) profiling revealed a highly active network involving multiple factors significantly upregulated upon cellular senescence but downregulated once cells are exposed to GSE treatment (Extended Data Fig. 1b). Further bioinformatics suggested that these molecules are functionally engaged in a group of biological processes including signal transduction, intercellular communication, energetic regulation, cell metabolism and inflammatory responses (Extended Data Fig. 1c). The majority of these downregulated genes were expressed as proteins delivered to extracellular space, located at endoplasmic reticulum or Golgi apparatus, consistent with the secreted nature of these molecules (Extended Data Fig. 1d).

Thus, GSE is a natural product that holds the potential to control the pro-inflammatory profile of senescent cells, namely the SASP, when used at a specified concentration. Although there were natural products that alternatively showed senolytic efficacy in cell-based assays (Fig. 1c), our subsequent study largely focused on GSE, as its geroprotective capacity appeared striking.

### GSE emerges as a powerful senolytic agent when applied at high concentrations

Given the prominent efficacy of GSE in reducing the SASP expression, we next interrogated the potential of this natural product in killing senescent cells at higher concentrations. To this end, we measured the percentage of senescent cells after treatment with increasing concentrations of GSE. Data from SA-β-Gal staining indicated that senescent cells were not significantly eliminated until GSE concentration reached 0.75 μg/ml (Fig. 2a-b). The efficacy of GSE in killing senescent cells was further increased with increasing concentrations, which reached a threshold (20% cell survival) when GSE was applied at 3.75 μg/ml, and did not further augment even at higher concentrations such as 7.50 μg/ml (Fig. 2a-b).

**Fig. 2.**
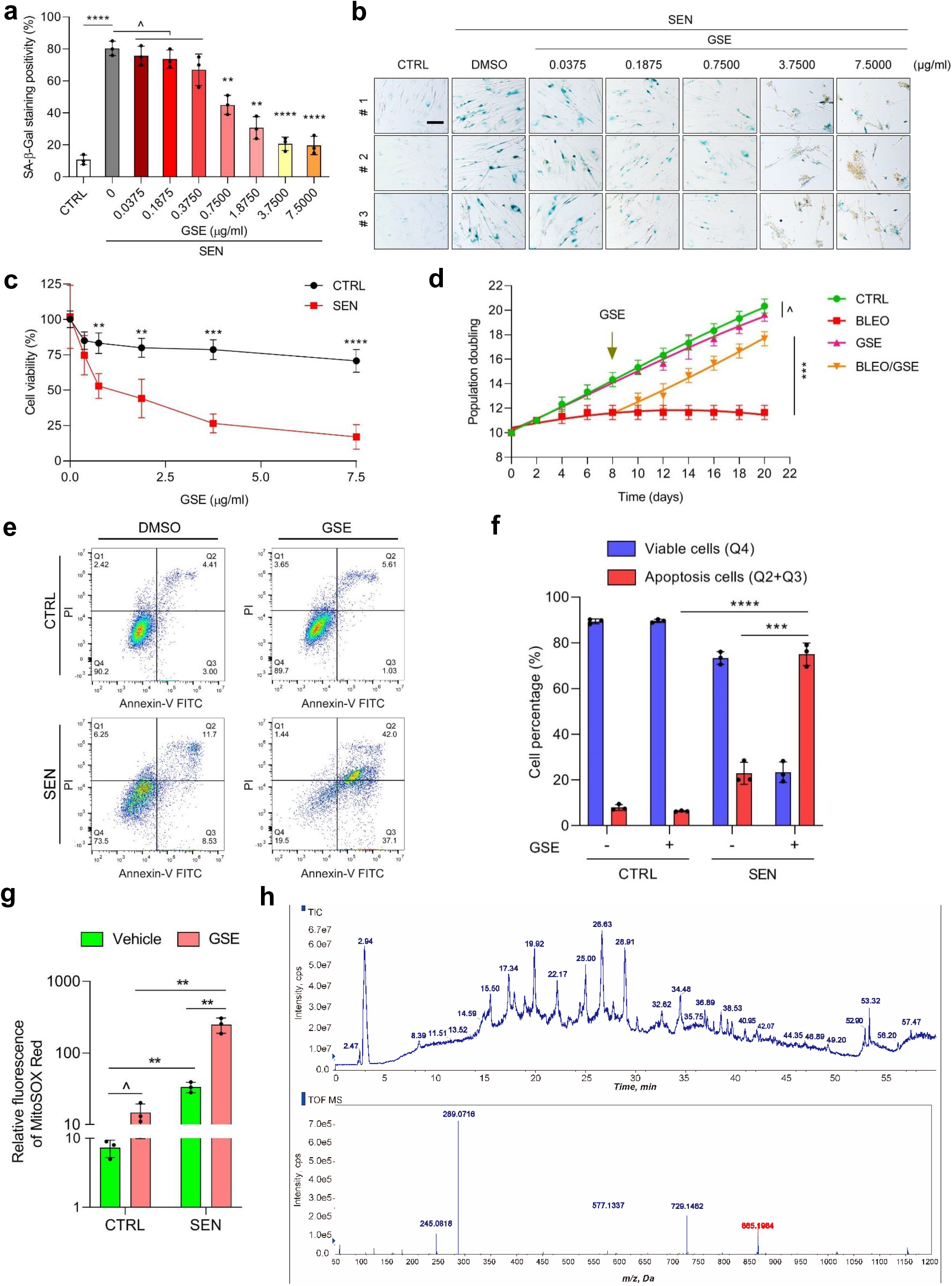
Characterization of the capacity of GSE in eliminating senescent cells. Quantification of senescent cell survival by SA-β-Gal positivity. GSE was applied to media at increasing concentrations. CTRL, control (proliferating) cells. SEN, senescent cells. (**b**) Representative images displaying SA-β-Gal staining results after PSC27 cells were treated by different concentrations of GSE. (**c**) Survival analysis of CTRL and SEN cells upon treatment by GSE. (**d**) Population doubling assessment of cells upon different treatments. GSE was applied on the 8^th^ day after beginning of experiments as indicated. (**e**) FACS measurement of CTRL and SEN cells after processing with an annexin V-FITC/PI kit and DAPI staining to determine apoptosis level. (**f**) Comparative quantification of the percentage of viable (Q4: PI− annexin V−) and apoptotic (Q2 and Q3: PI^+^/annexin V^+^ and PI^−^/annexin V+, respectively) cells in the CTRL or SEN populations treated with vehicle or GSE for 3 days (n = 3). (**g**) Measurement of the fluorescence signal of MitoSOX Red, a mitochondrial superoxide indicator, in PSC27 cells under different conditions. (**h**) High resolution mass spectra showing the total ion chromatogram (TIC) and base peak chromatogram (BPC) of GSE after performance of HPLC-ESI-QTOF-MS. Data in bar graphs and regression curves are shown as mean ± SD and representative of 3 biological replicates. ^, *P* > 0.05. *, *P* < 0.05. **, *P* < 0.01. ***, *P* < 0.001. ****, *P* < 0.0001.

We evaluated the potential of GSE as a senolytic agent with alternative methods. Cell viability assay showed that GSE induced significant death of senescent cells starting from the concentration of 0.75 μg/ml, in contrast to their proliferating counterparts (Fig. 2c). When GSE concentration was enhanced to 7.50 μg/ml, the percentage of surviving senescent cells declined to approximately 10%. However, there was no significant reduction in proliferating cells even at 15.00 μg/ml of GSE (data not shown), the highest concentration used in our cell assays, substantiating the selectivity and specificity of GSE against senescent cells, the major traits of senolytics as a distinct category of geroprotective agents.

We next examined the potential of population doubling (PD) of stromal cells after treatment with drug-delivered genotoxicity. For this purpose, we chose mesenchymal stem cells (MSC), which can resume colony-based proliferation even after exposure to certain extent of stressful treatment ^22^. In contrast to BLEO-damaged cells, which rapidly entered growth arrest after treatment, post-senescence (BLEO-induced) treatment with GSE significantly enhanced the PD capacity of these cells (Fig. 2d). However, treatment with GSE alone did not seem to affect the PD of proliferating cells, data further indicative of the selectivity of GSE between senescent cells and their normal counterparts.

Since GSE generates distinct effects against senescent cells, we analyzed the efficacy of GSE in inducing cell apoptosis. Flow cytometry demonstrated significantly reduced viability, while dramatically elevated apoptosis of senescent cells, but not proliferating cells (Fig. 2e-f). Mitochondrial dysfunction and metabolic changes are among the hallmarks of senescent cells and organismal aging, events causing oxidative stress and production of reactive oxygen species (ROS) such as the superoxide ^3, 23^. We used MitoSOX Red, a mitochondrial superoxide indicator ^24^, to probe intercellular changes, and found GSE1 promoted the generation of mitochondrial ROS in senescent, but not proliferating cells (Fig. 2g). Thus, our data consistently supported the prominent capacity of GSE in targeting senescent cells via induction of cell apoptosis and exacerbation of mitochondrial stress under *in vitro* conditions.

Grape seeds amount to 38%-52% on a dry matter basis of grapes, one of the most widely grown fruit crops worldwide, and constitute a handy source of antioxidant compounds ^25^. Since GSE exhibits remarkable senescence-targeting capacity, the phytochemical composition of GSE deserves thorough investigation. We applied high pressure liquid chromatography (HPLC) coupled to a quadrupole time-of-flight mass spectrometer (QTOF-MS) and equipped with an electrospray ionization (ESI) interface. The resultant data disclosed a series of compounds distributed in three major categories including phenolic acids, flavonoids (such as flavan-3-ol, procyanidins) and other compounds (Fig. 2h; Supplementary Table 1). Among them, a few components were identified as procyanidins and derivatives, which hold the potential to target mitochondrial proteins and ameliorate multiple chronic diseases ^26^. However, the major component(s) mediating the senolytic function of GSE remains largely unclear.

### The polyphenol component of GSE Procyanidin C1 is senolytic against senescent cells

Former studies reported the biological activities of grape seed procyanidins, such as reduction of oxidative damage, suppression of inflammation and induction of cancer cell apoptosis ^27–30^. Among individual compounds found in GSE, procyanidin C1 (PCC1) warrants special attention as it induces DNA damage, causes cell cycle arrest and increases expression of checkpoint kinases. Our preliminary analysis (total ions chromatogram, TIC) with HPLC-QTOF-MS of GSE, a mixture of phytochemical agents *per se*, suggested the presence of PCC1, as the profile of GSE at specific MS peaks is basically in line with the chromatogram profile of chemically pure PCC1 acquired from commercial sources (Fig. 2h and Extended Data Fig. 2a).

PCC1 decreases the level of Bcl-2, but increases the expression of Bax, activities of caspase 3 and 9 in cancer cells, thus holding anticancer property via induction of apoptosis ^31^. Thus, we next assessed the capacity and selectivity of PCC1 in eliminating senescent cells in culture. The data suggest that PCC1 is senolytic for senescent stromal cells starting from the concentration of 50 μM, wherein proliferating cells remain largely unaffected (Fig. 3a,b) (Supplementary Table 2). Although higher concentrations caused even lower survival rate of senescent cells, with a threshold approximately at 200 μM, the agent exhibited toxicity towards control cells only when used at 600 μM (data not shown). Data from time course assay indicated that PCC1 exerted apoptotic effects within 12 h after added to the media, and essentially reached a plateau 24 h post treatment of senescent cells (Fig. 3c). The senolytic nature of PCC1 was largely confirmed by data from senescent cells induced by replicative exhaustion (RS) or oncogene (*HRas^G12V^*) overexpression (OIS), stressors that result in consequences typically similar to those of therapy-induced senescence (TIS) (Fig. 3d and Extended Data Fig. 3a-d) (Supplementary Table 2). The results suggest that PCC1 selectively clears senescent human stromal cells induced by different stimuli in a dose-dependent manner, but without a significant effect on non-senescent cells when used at appropriate concentrations.

**Fig. 3.**
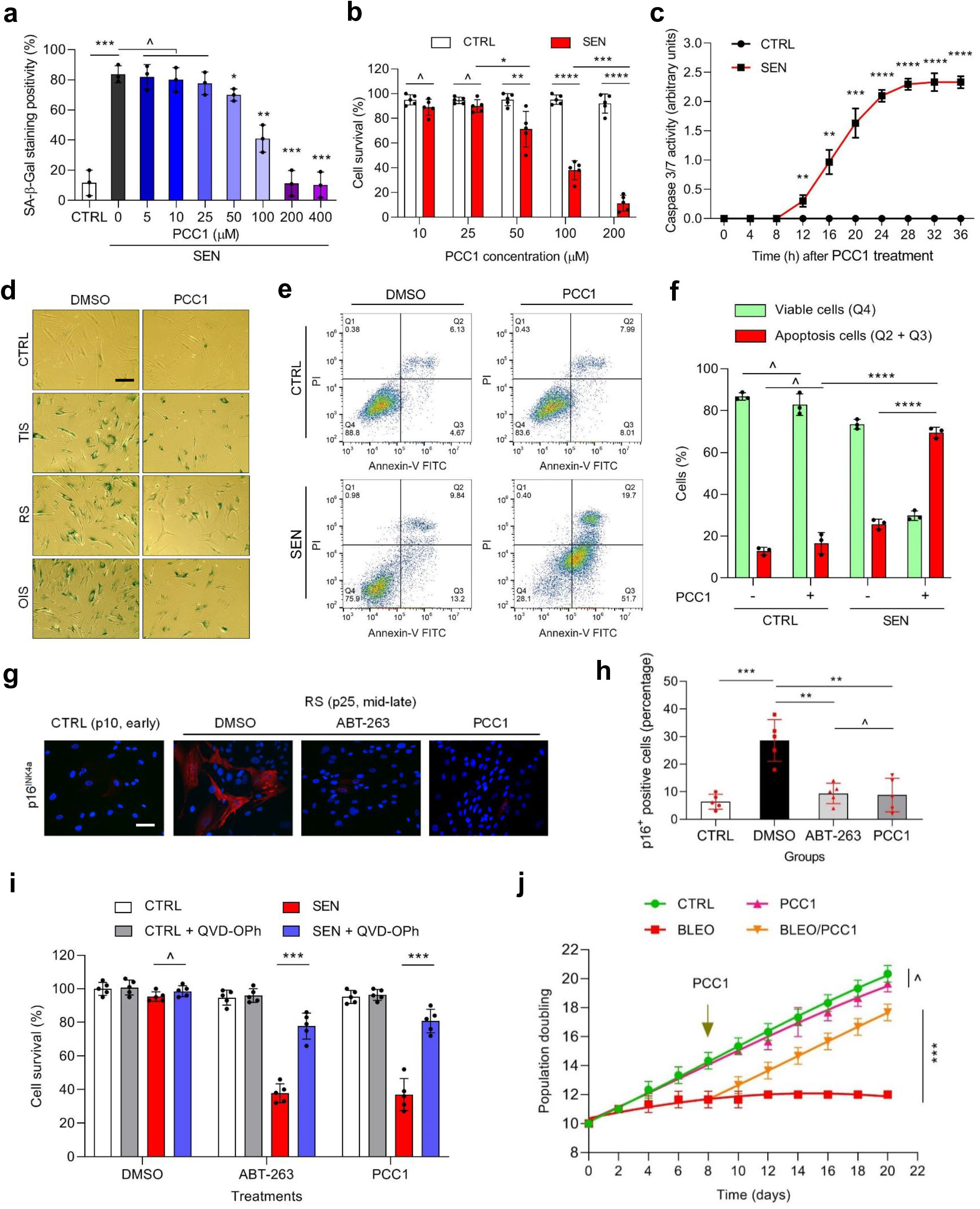
Characterization of the potential of PCC1 as a senolytic agent in targeting senescent cells. (**a**) Measurement of senescent cell survival by SA-β-Gal staining assay. Procyanidin C1 (PCC1) was applied at increasing concentrations. CTRL, proliferating. SEN, senescent. (**b**) Senolytic activity appraisal by calculating the percentage of surviving senescent cells induced by BLEO at increasing PCC1 concentrations. (**c**) Apoptotic assay of cells treated by PCC1 by examination of caspase 3/7 activity. (**d**) Representative images of SA-β-Gal staining upon treatment with vehicle or PCC1. CTRL, proliferating cells. TIS, therapy-induced senescence (by BLEO). RS, replicative senescence. OIS, oncogene-induced senescence (by oncogenic HRas^G12V^). Scale bar, 20 μm. (**e**) FACS measurement of CTRL and SEN cells after processing with an annexin V-FITC/PI kit and DAPI staining to determine apoptosis. (**f**) Comparative quantification of the percentage of viable (Q4: PI^−^/annexin V^−^) and apoptotic (Q2 and Q3: PI^+^/annexin V^+^ and PI^−^/annexin V^+^, respectively) cells in the CTRL or SEN populations treated with vehicle or GSE for 3 days (n = 3). (**g**) Immunofluorescence staining of PSC27 cells. RS was induced by consecutive proliferation, before cells were treated by PCC1. Red, p16^INK4a^. Cells at an early passage (p10) were used as a negative control. The senolytic agent ABT-263 was employed as a positive control. Scale bar, 20 μm. (**h**) Statistics of immunofluorescence staining positivity in assays described in (**g**). (**i**) Examination of PCC1-induced senolytic activity by pan-caspase inhibition (20 μM QVD-OPh). ABT-263 used as a positive senolytic control. (**j**) Population doubling assessment assay upon different treatments. PCC1 was applied on the 8^th^ day after the beginning of experiments as indicated. In experiments of **c**-**j**, PCC1 was used at 100 μm. Data are shown as mean ± SD and representative of 3 biological replicates. ^, *P* > 0.05. *, *P* < 0.05. **, *P* < 0.01. ***, *P* < 0.001. ****, *P* < 0.0001.

To experimentally expand and establish PCC1 efficacy across cell lineages, we treated human fetal lung fibroblasts (WI38), human umbilical vein endothelial cells (HUVECs) and human MSCs with PCC1, and found that senescent cells of these lineages exhibited similar susceptibility, which were selectively killed by PCC1, while their non-senescent counterparts sustained viability (Extended Data Fig. 3e-g) (Supplementary Table 3). Together, the data consistently support that PCC1 selectively eliminated senescent cells in a cell type- and senescence trigger-independent manner.

Next we applied flow cytometry to sort cells, and the resulting data showed markedly elevated apoptosis of senescent cells, while proliferating cells remained largely unaffected by PCC1 (Fig. 3e-f). Together, several lines of data suggested the substantial capacity of PCC1 in targeting senescent cells via induction of cell apoptosis, which can be considered as a novel senolytic agent derived from natural sources.

We further sought to visualize the real effect of PCC1 in depleting senescent cells exposed to PCC1 in media. As p16^INK4a^ is a widely used *in vitro* and *in vivo* marker of senescent cells, we examined its expression in stromal cells that experience replicative exhaustion. PCC1 remarkably removed p16-positive senescent cells, which only appeared in late passage PSC27 populations, with an efficacy largely resembling that of ABT263, a well-established synthetic senolytic agent originally designed to target cancer cells but later found to have the capacity of selectively killing senescent cells across multiple organ types ^18, 32–34^ (Fig. 3g-h).

To substantiate that PCC1-caused elimination of senescent cells is mainly through induction of apoptosis, rather than other forms of programmed cell death, we used the pan-caspase apoptosis inhibitor, QVD-OPh (quinolyl-valyl-O-methylaspartyl-[-2,6-difluorophenoxy]-methylketone) to treat cells. Although PCC1 significantly reduced the survival of senescent cells, the tendency was basically reversed by QVD-OPh, confirming caspase-dependent induction of apoptosis, a senolytic feature shared between PCC1 and ABT263 (Fig. 3i). Further analysis with chemical inhibitors against ferroptosis, pyroptosis or necroptosis basically excluded the possibility of PCC1-induced cell death through these mechanisms, while substantiating apoptosis as the major strategy of PCC-mediated senescent cell depletion (Extended Data Fig. 3h). In addition, population doubling assays further showed that PCC1 can resume the consecutive proliferating potential of MSC cells, which even exhibited a tendency to approach control cells despite initial senescence induction by BLEO-delivered genotoxicity in culture (Fig. 3j).

### PCC1 activates a pro-apoptotic signaling network to drive apoptosis of senescent cells

Given the prominent efficacy of PCC1 in selectively inducing senescent cell death, we interrogated the underlying mechanism(s). PCC1 belongs to the superfamily of flavonoids, which can scavenge free radicals, chelate metals, and reduce hydroperoxide formation, antioxidant properties attributable to the functional group “-OH” in the structure and its position on the ring of the flavonoid molecule ^25^. The antioxidant capacity of procyanidins is, in part, governed by their degree of polymerization, while PCC1 is a trimer procyanidin epicatechin by nature (Fig. 4a).

**Fig. 4.**
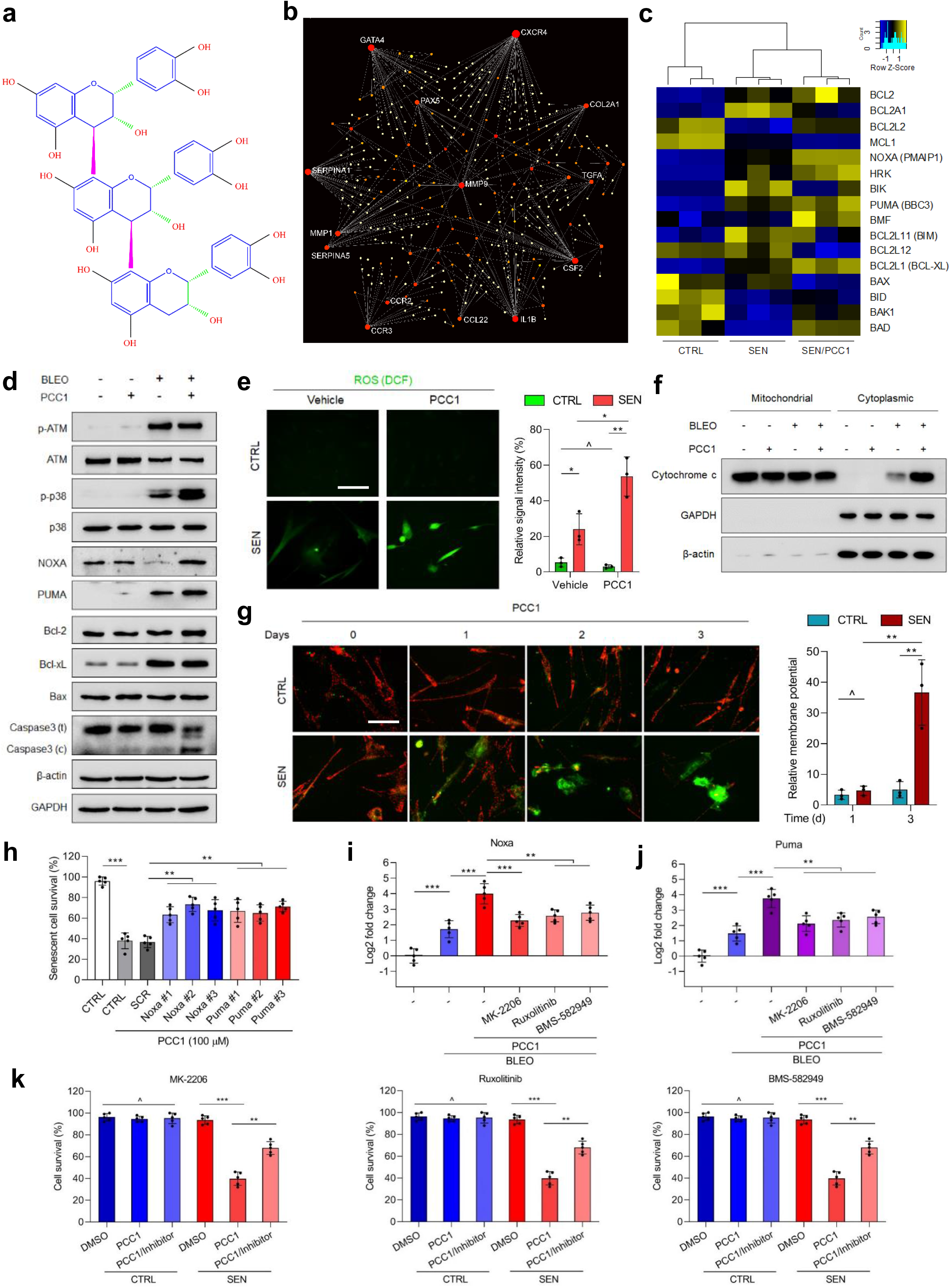
PCC1 induces apoptosis of senescent cells by engaging the pro-apoptotic pathways. (**a**) Chemical structure of the trimer epicatechin PCC1. (**b**) NetworkAnalyst map presentation of protein-protein interactions of typical SASP-associated factors significantly upregulated in senescent cells but downregulated by PCC1. (**c**) Heatmap showing the differential expression of Bcl-2 family genes in CTRL, SEN cells and PCC1-treated SEN cells. (**d**) Immunoblot analysis of PSC27 cells exposed to treatment by different agents. Expression of pro-apoptotic and anti-apoptotic factors was examined. Caspase (t), caspase total. Caspase (c), caspase cleaved. β-actin and GAPDH, loading controls. (**e**) Measurement of reactive oxygen species (ROS) in stromal cells with 2’-7’-dichlorodihydrofluorescein diacetate (DCFH-DA), a cell permeable fluorescent probe that is sensitive to changes in cellular redox state. Left, representative images. Right, statistics. (**f**) Immunoblot examination of cytochrome c in PSC27 cells exposed to different treatments. Distribution of cytochrome c between mitochondria and the cytoplasm was profiled via mitochondria isolation from cytosol supernatants. (**g**) PSC27 cells were subject to JC-1 staining, a fluorescent probe indicative of mitochondrial membrane potential. Green fluorescence indicates JC-1 monomers, which appear in the cytosol after mitochondrial membrane depolarization and indicate early stage apoptosis. Red fluorescence images JC-1 aggregation and resides on intact mitochondria. Left, representative images. Right, statistics. (**h**) PSC27 cells were infected with 3 different shRNAs targeting Noxa or Puma, before exposure to BLEO for senescence induction. Seven days later cells were treated by PCC1 (100 μM) for a 3-d period to induce apoptosis. SCR, scramble shRNA as non-specific control. (**i**) Noxa expression in PSC27 was determined using qRT–qPCR. Cells were treated by BLEO to induce senescence, before exposue to 100 μM PCC1 for 3 d, in the absence or presence of 10 μM MK2206, 10 μM Ruxolitinib, or 20 nM BMS-582949 to inhibit the activity of Akt, JAK1/2 or p38MAPK, respectively. (**j**) A similar set of qRT–qPCR expression assays for Puma in conditions described in (**i**). (**k**) Measurement of the viability of CTRL and SEN cells treated by PCC1 in the absence or presence of MK2206, Ruxolitinib or BMS-582949 against the enzymatic activity of Akt, JAK1/2 or p38MAPK, respectively. Statistical significance of **i**-**k** was calculated using Student’s t-test or one-way ANOVA (Dunnett’s test). Data of all bar graphs are shown as mean ± SD and representative of 3 biological replicates. ^, *P* > 0.05. *, *P* < 0.05. **, *P* < 0.01. ***, *P* < 0.001. ****, *P* < 0.0001.

We first analyzed the impact of PCC1 on the transcriptome-wide expression of senescent cells. Bioinformatics showed that 4406 genes were significantly upregulated, and 2766 genes downregulated in stromal cells upon PCC1 treatment (Extended Data Fig. 4a). We observed a large array of SASP factors, which were markedly upregulated upon cellular senescence but substantially downregulated when senescent cells were exposed to PCC1 (Extended Data Fig. 4b). GSEA data showed that both SASP signature and NF-κB signature were remarkably suppressed by PCC1 (Extended Data Fig. 4c-d). We further noticed multiple mutual interactions between these ‘senescent-up and PCC1 down’ factors appearing on the top list of differentially expressed genes, most of which are typical secreted factors (Fig. 4b).

To understand the selectivity of PCC1 towards senescent cells, we further assessed the transcriptomic expression profile and noticed that PCC1 induced expression changes of a few Bcl-2 superfamily members (Fig. 4c). Of note, PCC1-dependent upregulation/activation of p38 was observed, with caspase 3 cleavage occurring in these cells (Fig. 4d). Although Bcl-xL expression was elevated in senescent cells relative to their proliferating controls, PCC1 did not further enhance its protein level. The other two Bcl-2 factors, namely Bcl-2 and Bax, remained largely unchanged between different conditions. While Noxa and Puma exhibited different expression pattern upon cellular senescence, PCC1 treatment resulted in the upregulation of both factors (Fig. 4d).

As procyanidins increase cell viability, decrease ROS production and restrain oxidative stress in mammalian cells ^35, 36^, we next interrogated whether similar or antioxidant effects can be traced in senescent cells exposed to PCC1. Surprisingly, we observed the opposite side, as senescent PSC27 cells displayed elevated ROS levels when treated by PCC1, making a sharp contrast to their proliferating counterparts (Fig. 4e and Extended Data Fig. 4e, signals of DCFH-DA probe). Indeed, PCC1 exposure caused enhanced cytochrome c release from mitochondria to their surrounding cytoplasmic space, as evidenced by signals detected by immunoblots (Fig. 4f). Further analysis indicated that mitochondrial membrane potential (Δψm) was significantly elevated in senescent cells, while proliferating cells remained basically unaffected in the presence of PCC1, as indicated by the profile of JC-1 probe signals (Fig. 4g). Thus, PCC1 promotes ROS generation, cytochrome c release and Δψ_m_ disturbance in senescent cells, events inherently associated with apoptosis.

Further experimental assays involving knockdown of Bcl-2 pro-apoptotic factors suggested that Noxa and Puma, two molecules upregulated upon PCC1 treatment (Fig. 4d), mediated the senolytic actions of PCC1 (Fig. 4h and Extended Data Fig. 4f). Treatment with chemical inhibitors of Akt, JAK1/2, and p38 signaling supported involvement of these individual pathways in expression of Noxa and Puma and senescent cell apoptosis upon PCC1 treatment (Fig. 4i-k). Therefore, our data suggest that senescent cells are subject to PCC1- induced apoptosis, a process partially mediated by Noxa and Puma and characterized with remarkable mitochondrial dysfunction and elevated free-radical generation.

### Therapeutically targeting senescent cells with PCC1 promotes tumor regression and minimizes chemoresistance *in vivo*

Given the capacity and selectivity of PCC1 in eliminating senescent cells *in vitro*, we next interrogated whether this agent can be exploited to intervene age-related pathologies *in vivo*. Cancer represents one of the leading chronic diseases that severely threaten human lifespan and compromise healthspan. In clinical oncology, drug resistance limits the efficacy of most anticancer treatments, while senescent cells frequently contribute to therapeutic resistance via development of an *in vivo* SASP in the drug-damaged TME ^15, 16, 37^. However, the feasibility of depleting senescent cells from primary tumors to promote cancer treatment index remains largely underexplored.

First, we built tissue recombinants by admixing PSC27 with PC3, the latter a typical prostate cancer (PCa) cell line of high malignancy, at a pre-optimized ratio (1:4). The cells were subcutaneously implanted to the hind flank of experimental mice with non-obese diabetes and severe combined immunodeficiency (NOD/SCID). Animals were measured for tumor size at the end of an 8-week period, with tissues acquired for pathological appraisal. Compared with tumors comprising PC3 cancer cells and naïve PSC27 stromal cells, xenografts composed of PC3 cells and senescent PSC27 cells exhibited significantly increased volume (*P* < 0.001), basically confirming the promoting effects of senescent cells in tumor progression (Extended Data Fig. 5a).

To closely mimick clinical conditions, we experimentally designed a preclinical regimen incorporating genotoxic therapeutics and/or senolytics (Fig. 5a). Two weeks after subcutaneous implantation when stable uptake of tumors *in vivo* was observed, a single dose of MIT or placebo was delivered to animals at the 1^st^ day of 3^rd^, 5^th^ and 7^th^ week until end of the 8-week regimen and (Extended Data Fig. 5b). Contrasting to placebo-treated group, MIT administration caused remarkably delayed tumor growth, validating the efficacy of MIT as a chemotherapeutic agent (44.0% reduction of tumor size, *P* < 0.0001) (Fig. 5b). Notably, although administration of PCC1 itself did not cause tumor shrinkage, treatment of MIT followed by PCC1 delivery remarkably enhanced tumor regression (55.2% reduction of tumor size as compared with MIT only, *P* < 0.001; 74.9% reduction of tumor volume as compared with placebo-treated, *P* < 0.0001) (Fig. 5b).

**Fig. 5.**
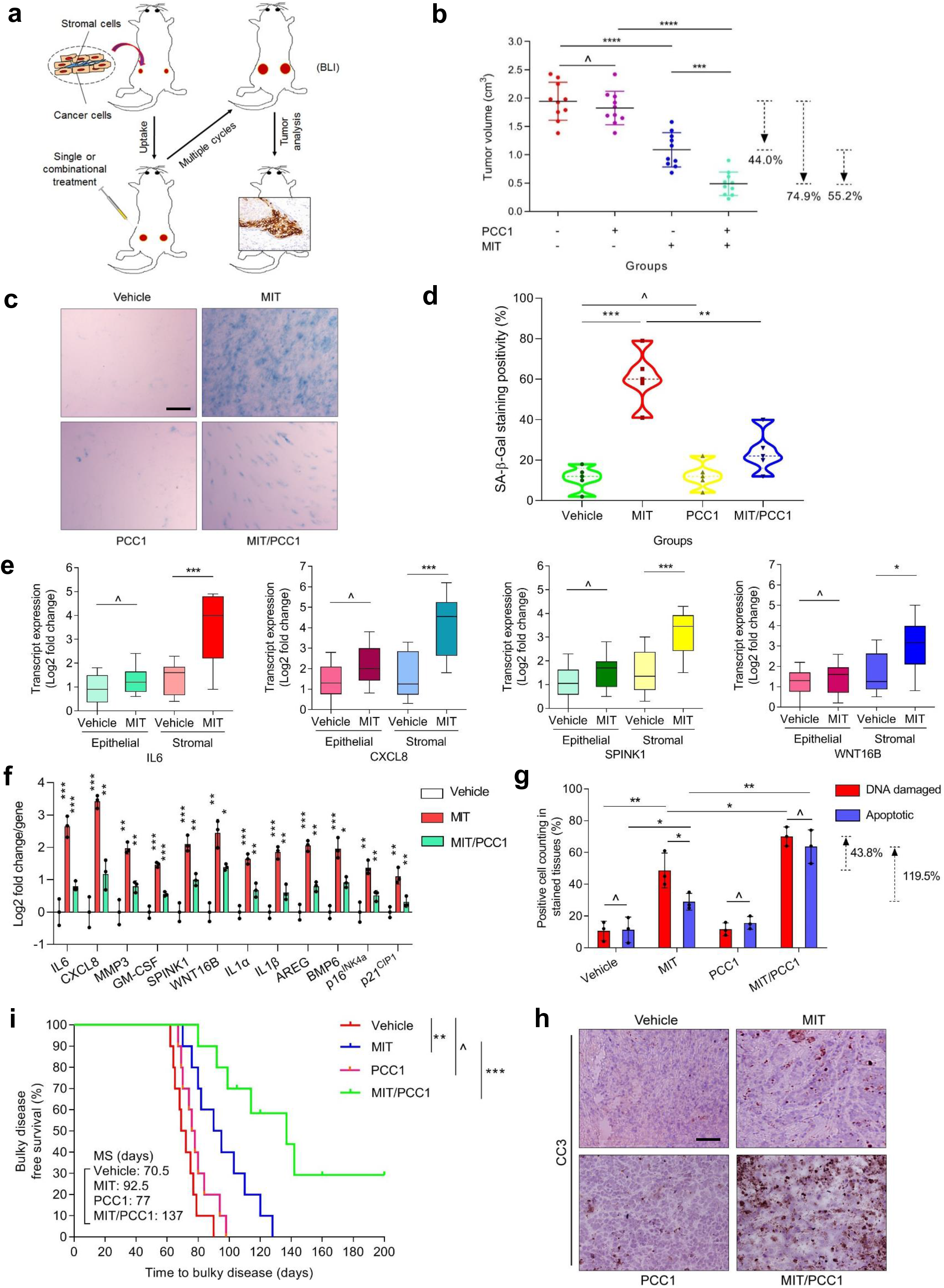
Senolytically targeting senescent cells in the damaged TME diminishes cancer resistance conferred by the SASP. (**a**) Illustrative diagram for a preclinical regimen. Two weeks after subcutaneous implantation and *in vivo* uptake of tissue recombinants, NOD/SCID mice received either single (mono) or combinational (dual) agents administered as metronomic treatments composed of several cycles. (**b**) Statistical profiling of tumor end volumes. PC3 cells were xenografted alone or together with PSC27 cells to the hind flank of animals. The chemotherapeutic drug MIT was administered to induce tumor regression, alone or together with senolytic PCC1. (**c**) Comparative imaging of *in vivo* senescence of tumor tissues assayed by SA-β-gal staining. Tumors were freshly dissected upon animal sacrifice and processed as frozen sections before tissue staining. Scale bars, 200 μm. (**d**) Violin plots that show comparative statistics of SA-β-gal staining positivity in tumor tissues. (**e**) Transcript assay of the *in vivo* expression of several canonical SASP factors (including IL6/CXCL8/SPINK1/WNT16B) in stromal cells isolated from tumors. Tissues from animals xenografted with both stromal and cancer cells were subject to LCM isolation, total RNA preparation and expression assays. (**f**) Quantitative assessment of SASP transcripts in stromal cells isolated from tumor tissues of animals subject to different treatments. Signals *per* factor were normalized to the vehicle-treated group. (**g**) Statistical measurement of DNA-damaged and apoptotic cells in the biospecimens collected as described in (**a**,**b**). Values are presented as percentage of cells positively stained by IHC with antibodies against γ-H2AX or caspase 3 (cleaved). (**h**) Representative IHC images of caspase 3 (cleaved, CC3) in tumors at the end of therapeutic regimes. Biopsies of vehicle-treated animals served as negative controls for MIT-treated mice. Scale bars, 100 μ (**i**) Survival analysis of mice sacrificed upon development of advanced bulky diseases. Survival duration was calculated from the time of tissue recombinant injection until the day of death. Data analyzed by log-rank (Mantel-Cox) test. Data of all bar graphs are shown as mean ± SD and representative of 3 biological replicates. ^, *P* > 0.05. *, *P* < 0.05. **, *P* < 0.01. ***, *P* < 0.001. ****, *P* < 0.0001.

We next reasoned whether cellular senescence occurs in the tumor foci of these animals. Not surprisingly, MIT administration induced appearance of a large number of senescent cells in tumor tissues. However, delivery of PCC1 to these chemo-treated animals essentially depleted the majority of senescent cells (Fig. 5c,d). Laser capture microdissection (LCM) followed by transcript assays indicated significantly elevated expression of SASP factors including IL6, CXCL8, SPINK1, WNT16B, GM-CSF, MMP3, IL1α, a tendency accompanied by upregulation of the senescence marker p16^INK4a^ in chemo-treated animals (Fig. 5e and Extended Data Fig. 5c). Intriguingly, these changes were mainly observed in stromal cells, rather than neighboring cancer counterparts, implying the possibility of repopulation of residual cancer cells, which frequently develop acquired resistance in the treatment-damaged TME. However, upon administration with PCC1, these changes were largely reversed, as suggested by transcript assays (Fig. 5f).

To investigate the mechanism directly responsible for the SASP expression in MIT-treated mice and reversal of such a senescence-associated pattern, we dissected tumors from animals treated by these two agents 7 days after the first dose of GSE delivery, a timepoint prior to the development of resistant colonies. In contrast to the placebo, MIT administration caused dramatically increased DNA damage and apoptosis. Although PCC1 alone did not induce DNA damage or apoptosis, the chemotherapeutic agent MIT did (Fig. 5g). However, when MIT-treated animals were administered with PCC1, the indices of both DNA damage and apoptosis were significantly augmented, implying an enhanced cytotoxicity in these chemo/senolytics-treated animals. As supporting evidence, there was elevated caspase 3 cleavage, a typical hallmark of cell apoptosis, when PCC1 was therapeutically applied to these MIT-treated mice (Fig. 5h).

We next evaluated the consequence of tumor progression by comparing the survival of different animal groups in a time-extended manner. In this preclinical cohort, animals were monitored for tumor growth, with bulky disease considered arising once the tumor burden was prominent (size ≥ 2000 mm3), an approach employed for certain situations ^14, 38^. Mice receiving MIT/PCC1 combinatorial treatment showed the most prolonged median survival, gaining a minimally 48.1% longer survival as compared with the group treated by MIT only (Fig. 5i, green *vs* blue). However, PCC1 treatment alone did not provide significant benefit, with only marginal survival extension conferred. The data suggest that PCC1 administration along neither changes tumor growth nor promotes animal survival, while MIT/PCC1 co-treatment can significantly improve both parameters.

Of note, treatments performed in these studies appeared to be well tolerated by animals, as no significant perturbations in urea, creatinine, liver enzymes, or body weight were observed (Extended Data Fig. 5d,e). More importantly, the chemo and geroprotective agents administered at doses designed in this study did not significantly interfere with the integrity of immune system and tissue homeostasis of critical organs even in immunocompetent mice (Extended Data Fig. 6a-c). These results consistently support that an anti-aging agent combined with conventional chemotherapy has the potential to enhance tumor response without causing severe systemic toxicity.

### Eliminating senescent cells prevents and alleviates physical dysfunction induced by senescent cells

Previously studies disclosed that even a small number of senescent cells can induce physical dysfunction in young animals ^39^. We asked whether PCC1 selectively kills senescent cells *in vivo*, thereby preventing physical dysfunction. To address this, we performed parallel implantation of control and senescent mouse stromal cells constitutively expressing luciferase (LUC^+^) subcutaneously into syngeneous WT mice. Animals were treated immediately after implantation with PCC1 or vehicle (V) for 7 days (Fig. 6a). We found luminescence signal intensities significantly lower in senescent cell-implanted mice treated with PCC1 than those in V-treated littermates, although no difference was observed following treatment of mice transplanted with LUC^+^ control cells (Fig. 6b,c), substantiating the senolytic efficacy of PCC1 *in vivo*.

**Fig. 6.**
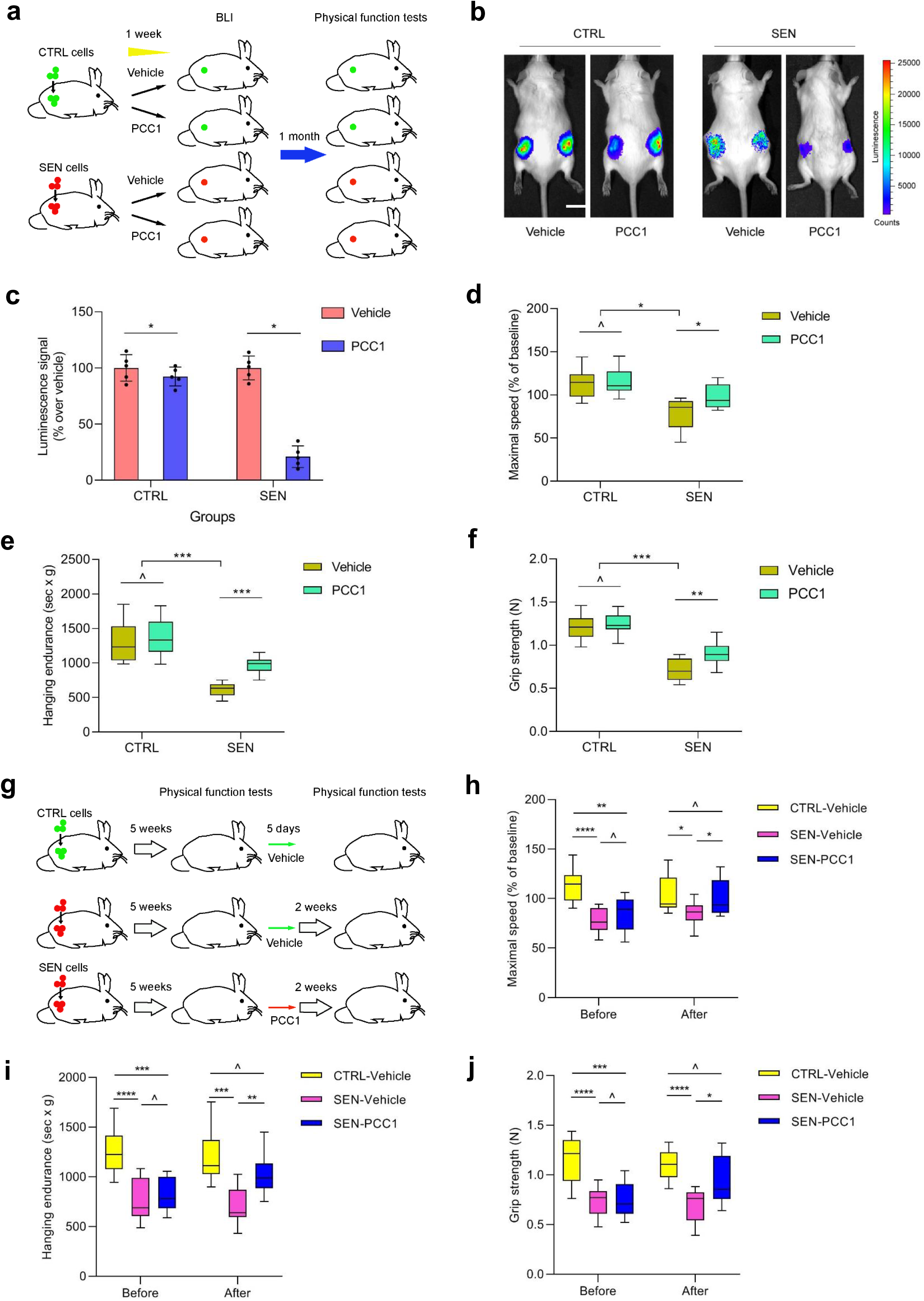
Elimination of senescent cells by PCC1 prevents physical dysfunction and alleviate pathological symptoms. (**a**) Schematic presentation of experimental procedure for cell transplantation and physical function tests of 5-month-old male C57BL/6J mice. (**b**) Representative images showing *in vivo* luciferase (LUC) activity 2 days after the last treatment of mice. Scale bars, 20 mm. (**c**) Luminescence of transplanted cells as a percentage in relative to the average of signals in mice treated with vehicle. (**d-f**) Measurement of maximal walking speed (relative to baseline) (**d**), hanging endurance (**e**) and grip strength (**f**) of 5-month-old male C57BL/6J mice, with tests performed 1 month after the last treatment. (**g**) Schematic experimental design for transplantation and physical function measurements. (**h**-**j)** Measurement of maximal walking speed (relative to baseline) (**h**), hanging endurance (**i**) and grip strength (**j**) of 28-week-old male C57BL/6J mice (2 weeks post the last treatment). Data are shown as box-and-whisker plots, where a box extends from the 25th to 75th percentile with the median shown as a line in the middle, and whiskers indicate smallest and largest values. ^, *P* > 0.05. *, *P* < 0.05. **, *P* < 0.01. ***, *P* < 0.001. ****, *P* < 0.0001.

We next interrogated whether killing implanted senescent cells using PCC1 can attenuate pathological events, specifically physical dysfunction. Treating young animals with PCC1 upon senescent cell implantation for a week prevented the decline of maximal walking speed (via RotaRod), hanging endurance (via hanging test) and grip strength (via grip meter), changes observed within 1 month after vehicle-treatment of another group of mice carrying senescent cells, consistent with the potential of PCC1 in ameliorating physical dysfunction (Fig. 6d-f). PCC1 administration also avoided physical dysfunction that occurred in animals 5 weeks following senescent cell implantation (Fig. 6g). In senescent cell-harboring mice, a single 5-day course of PCC1 treatment improved physical function compared to V group (Fig. 6h-j). Of note, the improvement was detectable 2 weeks after PCC1 treatment and lasted for even several months (Extended Data Fig. 7a,b). At these two time points of PCC1 administration (immediately *vs* 5 weeks after senescent cell implantation), the beneficial effects of PCC1 seem to be comparable. The data suggest that the time of administration of PCC1 may be flexible, implying its potential clinical potency. As plant seed-derived procyanidins usually have elimination half-lives of <12 hours ^40, 41^, such a sustained improvement in physical function following a single course of PCC1 treatment circumvents the need of continuous treatment with the senolytic agent, further implying that the activity of PCC1 is sufficient to avert senescent cell-induced physical dysfunction.

### Senescent cell clearance alleviates physical dysfunction and prolongs late-life survival without increasing morbidity in aged mice

Senolytics are able to deplete senescent cells in diverse tissue and organ types in various pathophysiological situations, most of which are correlated with aging ^42^. To further examine the effect of PCC1 on senescent cells in organisms, we selected two independent animal models of *in vivo* senescence, including therapy-induced mice and naturally aging mice. First, we induced cellular senescence by exposing wild type mice to whole body irradiation (WBI) at a sub-lethal dose (5 Gy), a step followed by geroprotective treatment via administration of PCC1 or vehicle (Extended Data Fig. 8a). Of note, animals that had undergone WBI manifested abnormal body appearance such as markedly greyed hair, which, however, was largely reversed by PCC1 administration (Extended Data Fig. 8b,c). SA-β-Gal-positive senescent cells were readily induced *in vivo* of these animals, as evidenced by enhanced staining positivity in specimens such as cardiac and pulmonary tissues (Extended Data Fig. 8d,e). However, when we treated these organ-injured animals with PCC1 by intraperitoneal injection (i.p.), the percentage of SA-β-Gal-positive cells in dissected tissues was significantly reduced, contrasting with that in vehicle-treated mice in post-WBI stage (Extended Data Fig. 8f,g). RT-qPCR results revealed that PCC1 also decreased expression of senescence markers and a subset of typical SASP factors compared with vehicle treatment (Extended Data Fig. 8h). Taken together, the data suggest that PCC1 could effectively deplete SA-β-Gal-positive cells, control the SASP expression and minimize senescence levels under *in vivo* conditions.

We then assessed the impact of preclinical treatments on the physical parameters of mice. As expected, WBI significantly compromised the exercise capacity and muscle strength as measured by treadmill and grip strength assays in the vehicle group (Extended Data Fig. 8i,j). In a sharp contrast, PCC1 administration provided substantial benefits by markedly recovering these capacities. More importantly, PCC1 treatment showed a prominent tendency to increase the survival rate (Extended Data Fig. 8k). Our results support that PCC1-induced elimination of SA-β-Gal-positive senescent cells may be an effective strategy to alleviate senescence-related physical regression and reduce mortality in settings of premature aging triggered by environmental stressors such as cytotoxic therapy.

Given the substansive effects of PCC1 treatment on accelerated aging, we next sought to define the impact of senescent cells on physical function in naturally aging animals, which indeed more closely mimick the pathophysiological changes in aging process *per se*. For this purpose, we chose to treat 20-month-old non-implanted, wild type (WT) mice with vehicle or PCC1 intermittently over 4 months (Fig. 7a). Histological evaluation revealed significantly elevated percentage of SA-β-Gal-positive senescent cells in major organs such as kidney, liver, lung and prostate of aged animals, a tendency that was markedly countered when PCC1 was administered as a senolytic agent (Fig. 7b,c and Extended Data Fig. 9a-f). Results from physical test showed that PCC1 ameliorated physical dysfunction, by enhancing maximal walking speed, hanging endurance, grip strength, treadmill endurance, daily activity and beam balance performance of animals administered with PCC1 compared to those treated with vehicle (Fig. 7d-i), although body weight and food intake remained largely unchanged in PCC1-treated mice (Extended Data Fig. 9g,h). Notably, expression of the SASP was significantly reduced in tissues such as those from lung of aged mice treated with PCC1 compared with vehicle group (Fig. 7j), a pattern largely consistent with lower secretion of the SASP factors by human stromal tissues treated by PCC1 (Fig. 5f).

**Fig. 7.**
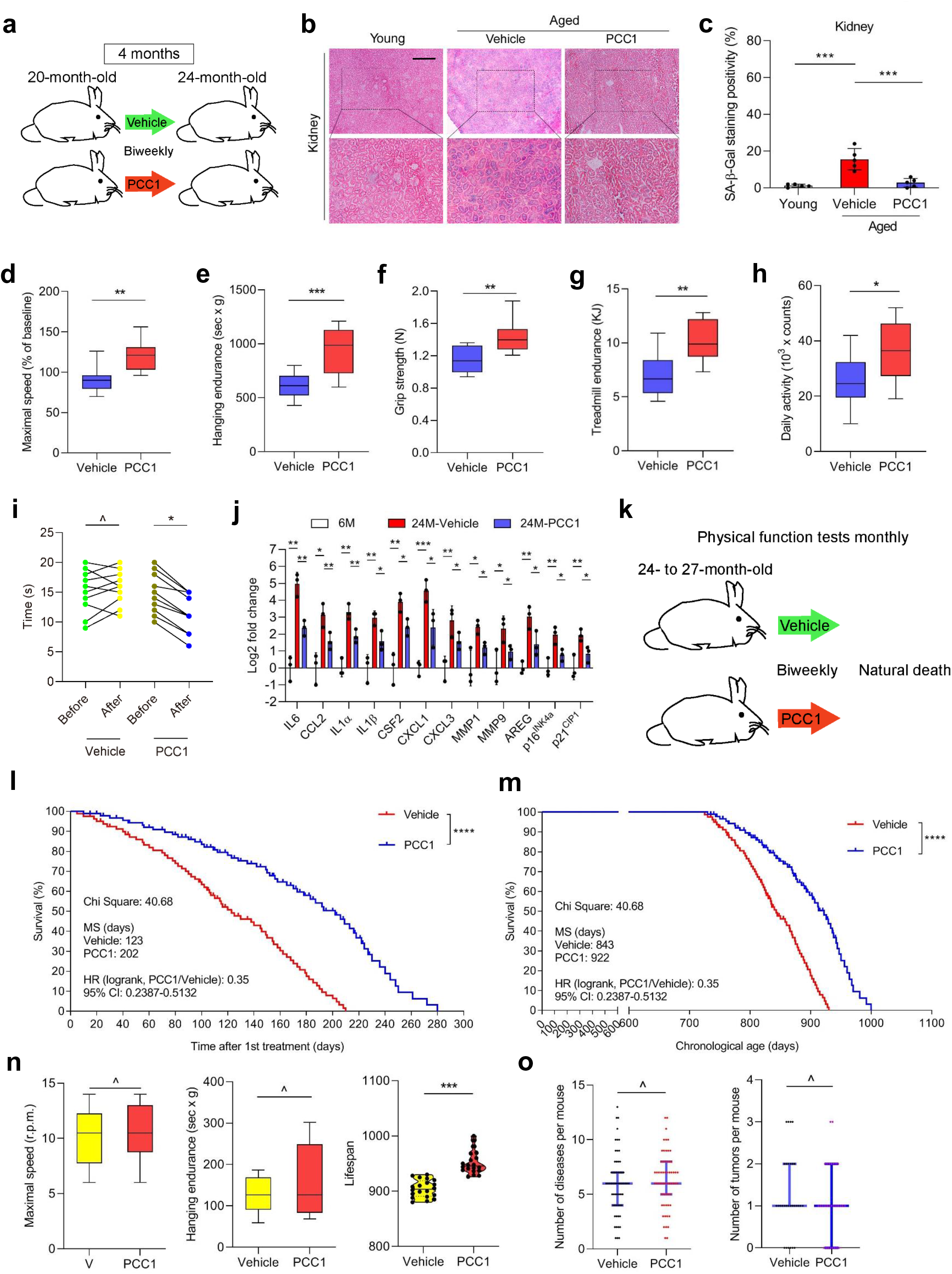
Intermittent administration of PCC1 extends both health- and lifespan of aged mice. (**a**) Schematic design for physical function measurements of 20-month-old male mice treated with PCC1 once per 2 weeks (biweekly) for 4 months. (**b**) Representative images of SA-β-Gal staining of kidneys from young and aged mice treated with vehicle or PCC1. Scale bar, 200 μm. (**c**) Quantification of SA-β-Gal staining in key tissues of animals as described in (**b**). (**d**-**h)** Quantification of maximal walking speed (relative to baseline) (**d**), hanging endurance (**e**), grip strength (**f**), treadmill endurance (**g**) and daily activity (**h**) of 20-month-old male C57BL/6J mice at 4 months after treatment was started. (**i**) Quantification of the time to cross the balance beam for vehicle-treated and PCC1-treated old mice. Data of before and after treatment *per* animal were connected to allow direct comparison of treatment effect. (**j**) Quantitative measurement of SASP factor expression at transcription level in lung tissues collected from 6-month-old untreated (6M), 24-month-old vehicle-treated (24M-Vehicle) and 24-month-old PCC1-treated mice (24M-PCC1). (**k**) Schematic design for lifespan analyses of experimental mice at 24-to-27 months of age. (**l**-**m**) Post-treatment survival (**l**) and whole-life survival (**m**) curves of C57BL/6J mice treated biweekly with PCC1 (*n* = 91; 48 males, 43 females) or vehicle (*n* = 80; 42 males, 38 females) starting at 24-27 months of age. Median survival is shown for all groups. (**n**) Maximal walking speed and hanging endurance averaged over the last 2 months of life, and lifespan for the longest living mice (top 20) in both groups. (**o**) Disease burden and tumor burden at death. For both sexes, *n* = 60 for V or PCC1. For males, *n* = 31 for PCC1, *n* = 33 for V. For females, *n* = 29 for PCC1, *n* = 27 for V. Data are displayed as box-and-whisker plots, where a box extends from the 25th to 75th percentile with the median shown as a line in the middle, and whiskers indicate smallest and largest values (**d**-**h**, **n**), or as mean ± SD (**i**). ^, *P* > 0.05. *, *P* < 0.05. **, *P* < 0.01. ***, *P* < 0.001. Unpaired two-tailed Student’s *t*-tests (**c**-**j**, **o**) and Cox proportional hazard regression model (**l**, **m**).

Further, to establish the potential of senescent cell elimination in extending the remaining lifespan of WT mice, we performed intermittent treatment with PCC1 beginning at very old age (Fig. 7k). Notably, mice receiving bi-weekly administration of PCC1 starting at 24-27 months of age (equivalent to age 75-90 years in humans) had 64.2% higher median post-treatment lifespan (or 9.4% higher overall lifespan) and lower mortality hazard (65.0%, *P* < 0.0001), compared to the vehicle group (Fig. 7l,m). These data indicate that PCC1 can significantly decrease the risk of age-associated mortality in old mice.

We next asked whether the reduced death rate in aged animals comes at the cost of an increased late-life morbidity, we measured physical function in experimental mice treated with PCC1 or vehicle monthly until death. Despite the longer remaining lifespan in PCC1-treated mice, physical function in the last 2 months of life was not significantly lower compared to vehicle-treated mice (Fig. 7n). Upon autopsy, the incidence of several age-related pathologies, tumor burden and cause of death, were not significantly different between PCC1- and vehicle-treated mice (Fig. 7o and Extended Data Fig. 10a,b). However, expression of the SASP was effectively reduced in solid organs, largely compatible with the decline of circulating levels of IL6, CSF2 and MCP1, representative SASP markers in the peripheral blood (Extended Data Fig. 10c-f). We also observed decreased expression of the SASP in peripheral blood CD3^+^ T cells (Extended Data Fig. 10g), a cell lineage that exhibits a robust increase of p16 expression during human aging ^43^. Furthermore, PCC1 reduced oxidative stress in liver tissues as evidenced by a decrease of lipid peroxidation product 4-hydroxynonenal (HNE) adducts and an increase in the ratio of reduced to oxidized glutathione (Extended Data Fig. 10h,i), consistent with the general properties of flavonoids which have antioxidant activity by counteracting free radicals and engaging the antioxidant defense system ^44, 45^.

Together, the senolytic agent PCC1, a phytochemical component derived from GSE (or alternatively, in less abundance, from natural products such as extracts of cinnamon, cacao, apple peel and pine bark), can reduce the burden of senescent and possibly other cells developing a pro-inflammatory phenotype and inherently dependent on pro-survival SCAP pathways, and increase post-treatment lifespan without causing elevated morbidity in mice. We hereby present proof-of-principle evidence that even when administered in late life, such a therapeutic modality holds prominent potential to remarkably delay aging, restrains age-related diseases and enhances health conditions, thus providing a new avenue to improve healthspan and lifespan in future geriatric medicine.

## Discussion

Aging is an essentially inevitable process that progressively causes functional decline in nearly all organisms. Cellular senescence, a state of permanent growth arrest, has recently emerged as both a hallmark and a driver of aging ^3, 46^. Senescent cells accumulate in aged tissues over time and contribute to an increasing list of pathologies ^47^. Clearance of senescent cells from progeroid or naturally aged mice extends healthspan, increases lifespan and restrains age-related disorders including but not limited to atherosclerosis, osteoarthritis and neurodegenerative diseases.^48–51^. Recent advances in age-related studies prompted a wave for search of drugs that can selectively kill senescent cells, particularly a new class of geroprotective agents termed senolytics. A handful of senolytics are reported, including dasatinib and quercetin, piperlongumine, HSP90 inhibitors and Bcl-2 family inhibitors such as ABT-263 (navitoclax) and ABT-737 ^11, 13, 18, 19, 52^. Among them, Bcl-2 inhibitors are the most widely used senolytics, although originally developed as therapies for lymphoma. ABT-737 targets Bcl-2, Bcl-_X_L and Bcl-w, but with low solubility and oral bioavailability. More effective for *in vivo* use, ABT-263 mainly inhibits Bcl-2 and Bcl-_X_L, whereas it frequently causes thrombocytopenia. Given the marked side-effects of most senolytic compounds, there is a need to identify new compounds of senolytic activity but with reduced cytotoxicity. In this study, we screened a drug library composed mainly of natural products, with an aim to identify novel agent(s) that can widely target senescent cells with optimal *in vivo* efficacy and safety. As a result, we identified PCC1, a phytochemical agent derived from natural sources, as a broad-spectrum senolytic compound.

Genetic and pharmacological strategies demonstrated an array of benefits of eliminating senescent cells to delay aging and control diseases. Cellular senescence can be triggered by a variety of stimuli ranging from oncogenic activation, genotoxic stress, to inflammatory response and replicative exhaustion. Several compounds are identified as broad-spectrum senolytics, while others are selective against only a certain type of senescent cells. Difference in specificity implies the individual choices of senolytics, which mainly depends on their intended clinical use. A recent study revealed ouabain, a natural compound belonging to the cardiac glycoside (CG) family, as a senolytic agent that can be used for both senescent cell elimination and cancer therapy, the latter implemented via a dual mechanism of action ^53^. In this work, we discovered PCC1 as another novel, natural and potent senolytic, which selectively and specifically induces apoptosis of senescent cells, but with limited cytotoxicity to proliferating cells ^54^. Mechanistically, PCC1 targets mitochondrial membrane potential (Δψm) and generates substantial free-radicals such as the ROS in senescent, rather than proliferating cells. Senescent cells tend to develop a depolarized plasma membrane and have increased concentrations of H+^54^, a feature that makes them more susceptible to the action of PCC1. These alterations are accompanied by upregulated expression of pro-apoptotic factors Noxa and Puma, events that together result in senescent cell apoptosis. The vulnerability of senescent cells to PCC1 can be exploited for therapeutic purposes, as evidenced by tumor eradication *in vivo* of experimental mice, a case involving the combination of a senogenic drug and a senolytic agent and achieving the optimal treatment effects.

Cellular senescence *per se* is a highly heterogeneous process depending on different cell origins and environmental stimuli ^55^. One of the key features of PCC1 is its ability to efficiently clear senescent cells in a wide spectrum of cell types and stressors, including replication, oncogene, irradiation and chemotherapy. In this study, we compared PCC1 with other reported senolytics on human stromal cells, fibroblasts, HUVECs and MSCs, several major cell types in the tissue microenvironment. As reported, ABT263 eliminates senescent HEFs and HUVECs, but shows little effect on human preadipocytes, ^12, 18^. Combined use of dasatinib and quercetin (‘D + Q’) can deplete all three types of senescent cells in a dose-dependent manner, but is toxic against proliferating cells ^11, 56, 57^. Fisetin, another natural flavonoid reported as a senolytic agent, displays modest effect on senescent HEFs and preadipocytes only at high concentrations ^20, 58^. In contrast, PCC1 holds potential to overcome these limitations, including cell-type dependency, high toxicity on non-senescent cells, and low efficiency on senescent cells. Thus, PCC1 has a superior senolytic activity with excellent specificity and efficiency on a wider range of cell types, and covers several major types of senescence inducers.

We found that PCC1 exerts apoptosis-inducing effect on senescent cells under *in vivo* conditions. PCC1 eliminated therapy-induced senescent cells effectively and reduced senescence markers in solid organs, highlighting its effectiveness *in vivo*. In this study, we also treated naturally aged mice with PCC1 and tested its effects on senescent cells, chronic inflammation and physical function. First, PCC1 depleted senescent cells in multiple tissues and decreased the SASP-associated signatures as shown by the GSEA analysis. Second, PCC1 could suppress expression of the SASP-associated genes in aged livers and kidneys and reduce chronic low-grade inflammation in the blood. Third, PCC1 ameliorated the impaired motor function, balance, exhausted exercise, muscle strength, and spontaneous exploration in aged mice. Most importantly, the performance of RotaRod and beam balance in the PCC1-treated group was improved compared with that in the initial pretreatment condition. Collectively, the phytochemical drug PCC1 targeting senescent cells in the tissue microenvironment generated remarkable biological effects in naturally aged mice.

Similar to chemically synthesized counterparts, naturally derived procyanidins manifest anti-inflammatory, anti-arthritic, anti-allergic and anti-cancer activities, scavenge oxygen free radicals and suppress radiation-induced peroxidation activity ^59, 60^. As an epicatechin trimer isolated from plant materials mainly grape seeds, PCC1 provides health benefits in chronic pathological conditions, many of which are age-related ^61^. However, insightful evaluation of the toxicological effects of PCC1 *in vivo* is critical to its clinical applications. Our data showed that high concentration (20□mg/kg) and high frequency PCC1 treatment had no apparent systemic toxicities, providing solid evidence for *in vivo* safety of this senolytic. Moreover, the short duration of PCC1 treatment helps avoid significant impairment in tissues and further minimizes its side effects. In summary, our study demonstrates the superiority and safety of a geroprotective strategy by selectively targeting senescent cells in aged or treatment-damaged tissues, with a broad spectrum of cell types. These findings together open a new avenue for extending healthspan and prolonging lifespan, and treating age-related pathologies, with a senolytic agent that is from natural sources and has prominent efficacy.

## Methods

### Cell lines, *in vitro* culture and lentivirus

Human primary prostate stromal cell line PSC27 was kindly provided by Dr. Peter Nelson and cultured in PSC complete medium as described previously ^15^. Human fetal lung fibroblast cell line WI38 and human umbilical vein endothelial cell (HUVEC) line were purchased from ATCC, and maintained in DMEM and F-12K Medium, respectively, as recommended by the provider. Mesenchymal stem cell (MSC) line was derived from human umbilical vein tissues and cultured in MSC complete media with 10 μg/ml recombinant human insulin as reported ^62^. All cell lines were tested negative for microbial contamination and routinely authenticated with STR assays.

Lentiviral particles were produced using Lipofectamine 2000 and a packaging kit (Thermal Scientific) based on manufacturer’s instructions. PSC27 infected with viruses encoding the puromycin resistance gene were selected using puromycin (1 μg/ml) for 3 days.

### Cell treatments and appraisal

Stromal cells were grown until 60%-80% confluent (CTRL) and treated with bleomycin (50 μg/ml, BLEO, MedChemExpress) for 12 h. After treatment, the cells were rinsed twice with PBS and allowed to stay for 7-10 d in media. Alternatively, cells were allowed to passage consecutively for replicative exhaustion as replicative senescence (RS). Cells were also lentivirally infected with a construct encoding full length HRAS^G12V^ and selected by puromycin (1μg/ml) for 3 days and allowed to stay for 7 days until senescence.

DNA-damage extent was evaluated by immunostaining for γH2AX or p53-BP1 foci by following a 4-category counting strategy as formerly reported ^15^. Random fields were chosen to show DDR foci, and quantified using CellProfiler (http://www.cellprofiler.org). For clonogenic assays, cells were seeded at 1 × 10^3^ cells/dish in 10mm dishes for 24 h before treated with chemicals. Cells were fixed with 2% paraformaldehyde 7-10 days post treatment, gently washed with PBS and stained instantly with 10% crystal violet prepared in 50% methanol. Excess dye was removed with PBS, with plates photographed. Colony formation were evaluated by quantifying the number of single colonies *per* dish.

### Vector construction

The pLKO.1-Puro plasmid was purchased from Addgene. Small hairpin RNAs were cloned for generation of knockdown constructs (shRNAs) (oligos listed in Supplementary Table 4).

### Cell viability assays

Senescent or control cells were seeded into wells of a 96-well plate before treated with vehicle (0.1% DMSO or PBS) or natural product library agents (listed in Supplementary Table 5) with 5,000 cells *per* well at three increasing concentrations (1 μg/ml, 5 μg/ml and 10 μg/ml for agents of unknown MW; or 1 μM, 5 μM and 10 μM for agents of known MW) for a consecutive period of 3 days. Cells were digested with 0.25% trypsin and 1 mM EDTA, and harvested in PBS containing 2% FBS, with survival measured by cell-based MTS assay (Promega). The top candidate agents were further screened by a 30-day treatment at more specified concentrations. MSC cells (passage 5-10) were seeded into 6-well plates at a density of 30,000 cells *per* well. Culture medium with different candidate agents was changed every other day. The population doubling potential was evaluated as the relative cell proliferative abilities of MSCs treated with each drug under culture conditions.

Apoptosis was assessed with an Annexin V-FITC cell apoptosis assay kit (Beyotime). Briefly, cells were sequentially incubated with Annexin V-FITC and propidium iodide (PI), before subjected to sorting by a BD LSR II flow cytometer (BD Biosciences). Viable cells (PI^−^ cells) were analyzed at a constant flow rate and calculated as a percentage of control cells treated with vehicle using the formula: percentage of control = (*N*drug/*N*c) × 100, where *N*drug and *N*c represent the absolute number of PI^−^ viable cells for drug-treated and vehicle-treated cells, respectively. Dose-response curves were established for each tested agent and the half-maximal inhibitory concentrations (IC50 values) were calculated by Probit analysis ^63^.

### SA-**β**-galactosidase staining, BrdU incorporation and apoptosis assays

SA-β-Gal staining assay was carried out with a SA-β-Gal staining kit (Beyotime) according to the manufacturer’s instruction. Alternatively, cells were incubated with X-Gal staining solution (1 mg/ml X-Gal, Thermo Scientific) with 5 mM K_3_[Fe(CN)_6_] and 5 mM K_4_[Fe(CN)_6_] for 12-16 h at 37°C. The percentage of SA-β-Gal positive cells was estimated by counting at least 100 cells per replicate sample facilitated by the “point picker” tool of ImageJ software (NIH). Flow SA-β-Gal staining assay was performed by flow cytometry using an ImaGene Green C12FDG lacZ gene expression kit from Molecular Probes (Life Technologies), according to the manufacturer’s instructions.

To examine the proliferation or senescence status of cells, a BrdU incorporation assay (10 μM, Yeasen) was performed for 16-18 h and cells were subject to immunofluorescence staining with an anti-BrdU antibody (1:400 dilution, Cell Signaling, cat no. 5292) before counterstained with DAPI and assessed by immunofluorescence and high content analysis microscopy. For crystal violet staining, the cells were seeded at low density on 6-well dishes and fixed at the end of the treatment with 0.5% glutaraldehyde (w/v). The plates were then stained with 0.2% crystal violet (w/v).

For apoptosis assays, target cells (proliferating or senescent) were plated in 12-well plates at 1.2 × 10^5^ cells/well. Twenty-four h after seeding, cells were treated with specific concentrations of geroprotective agents (ABT-263 at 1 μM, ABT-737 at 10 μM, Selleckchem; GSE at 0.75 μg/ml, or PCC1 *at 50* μM, TargetMol) for 72 h. The percentage of survival was determined based on quantification of remaining adherent cells using PrestoBlue reagent (A13262, Life Technologies) relative to control (DMSO-treated) cells.

To counteract apoptotic activity, a potent pan-caspase inhibitor QVD-OPh (MedChemExpress) was applied to culture media at concentration of 1 μM before addition of geroprotective agents.

### IncuCyte analysis

PSC27 cells were plated in 96-well dishes and induced to undergo senescence via BLEO treatment at 50 μg/ml. GSE and PCC1 were added at 0.75 μg/ml and 50 μM, respectively. The cell culture medium was supplemented with IncuCyte NucLight Rapid Red reagent for cell labelling (Essen Bioscience) and IncuCyte caspase-3/7 reagent for apoptosis (Essen Bioscience). Four images per well were collected every 4 h for 3 d.

### Mitochondrial isolation

Stromal cells cultured under regular conditions or experiencing drug treatments were subject to lysis and acquisition of mitochondria with a Cell Mitochondria Isolation Kit (Beyotime) by following the manufacturer’s protocol. Briefly, 1×10^8^ cells were harvested and washed twice with ice-cold PBS, with pellets resuspended with 600 μl ice-cold buffer A (20 mM HEPES [pH 7.5], 1.5 mM MgCl_2_, 10 mM KCl, 1 mM EDTA, 1 mM ethylene glycol bis (2-aminoethyl ether) tetraacetic acid, 1 mM dithiothreitol, 0.1 mM phenylmethanesulfonyl fluoride, and 1% protease inhibitor cocktail) for 10 mins and then added 200 μl 1 M sucrose to make the final concentration at 250 mM. Thereafter, cell suspension was passed five times through a 26-gauge needle fitted to a syringe quickly crushing cells to make at least 80% of the cells broken. Large plasma membrane pieces, nuclei, and unbroken cells were removed by centrifuging at 1,000 × *g* at 4°C for 10 mins. The supernatant was subjected to 10,000 × *g* at 4°C for 20 mins. The pellet (mitochondria) was rinsed with the above buffer A containing sucrose. The supernatant was re-centrifuged at 100,000 × *g* at 4°C for 1 h to generate cytosol. The pellet of mitochondria and cytosol was kept at −80°C for later use.

### Intracellular reactive oxygen species measurement

The intracellular reactive oxygen species (ROS) level was determined using a ROS Assay Kit (Beyotime) which uses dichloro-dihydro-fluorescein diacetate (DCFH-DA) as a probe. Briefly, cells were cultured in 6-well plates for 24 h at 37°C and were then washed twice with serum-free medium. Medium containing 10 µM DCFH-DA was added. Cells were then incubated for 20 min at 37°C, with light avoided during incubation. After incubation, the cells were washed thrice with serum-free medium, then observed and photographed using a fluorescence microscope (Nikon). The fluorescence intensity was quantitatively measured using ImageJ software (v1.51, NIH).

### Mitochondrial membrane potential (ΔΨm) analysis

The JC-1 probe was used to measure apoptosis via mitochondrial depolarization. Briefly, cells (80% confluent) were cultured in 96-well plates after treatment with 50 µM NPPB at 37°C for 24 h. Carbonyl cyanide 3-chlorophenylhydrazone (CCCP) was used as control, which is a chemical inhibitor of oxidative phosphorylation and decreases the potential of mitochondria membrane. The cells were washed thrice with PBS, and 50 µl medium and 50 µl JC-1 working solution were added. The cells were incubated at 37°C for 20 min and washed thrice with pre-cooled JC-1 solution. The ΔΨm was determined by measuring the fluorescence intensity of red and green fluorescence using a fluorescence microscope (Nikon), with green fluorescence indicative of JC-1 monomers. JC-1 monomers appears in cytosol after mitochondrial membrane depolarization, indicating early stage apoptosis. Red fluorescence suggests JC-1 aggregation and is located on the mitochondria. Mitochondrial depolarization implies apoptosis, as reflected by an increase in the green/red fluorescence intensity ratio. Fluorescence intensity was measured using ImageJ (NIH).

### Immunoblotting analysis

Cells or tissues were homogenized in lysis buffer (Cell Signaling) supplemented with protease inhibitors (Sigma), with resultant lysates cleared by centrifugation (16,000 × *g* for 10 min at 4 °C). Total protein content was determined using Coomassie Brilliant Blue G-250 Plus reagents (Pierce). Proteins were loaded to each lane on a 4-20% gradient SDS/PAGE gel and transferred to nitrocellulose membranes (Bio-Rad). Signals were developed with HRP-conjugated secondary antibodies and SuperSignal West Pico Chemiluminescent Substrate (Pierce). Antibodies were purchased from individual commercial sources (Supplementary Table 7).

### Immunofluorescence staining

Cells were fixed with 4% (v/v) paraformaldehyde (PFA) for 10 min and permeabilized in PBS containing 0.25% Triton X-100 for 10 min. After PBS washes, cells were blocked with PBS and Tween-20 (PBST) containing 5% BSA for 30 min. Cells were incubated with primary antibodies in 5% BSA in PBST overnight at 4 °C, then with the Alexa Fluor 488 or 594-conjugated secondary fluorescent antibodies (Life Technologies) in the same buffer for 1 h in dark. DAPI (1 μg/ml, Life Technologies) was used to counterstain nuclei. Slides were mounted with Vectashield medium. Fluorescence imaging was performed on a fluorescence microscope (Nikon Eclipse Ti S). Captured images were analyzed and processed with the Nikon DS-Ri2 fluorescence workstation and processed with NIS-Elements F4.30.01. Alternatively, a confocal microscope (Zeiss LSM 780) was applied to acquire confocal images (antibodies used for this study listed in Supplementary Table 7).

### Histology and immunohistochemistry

Mouse tissue specimens were fixed overnight in 10% neutral-buffered formalin and processed for paraffin embedding. Standard staining with hematoxylin/eosin was performed on sections of 5-8 μm thickness processed from each specimen block. For immunohistochemistry, tissue sections were de-paraffinized and incubated in citrate buffer at 95 °C for 40 min for antigen retrieval before incubated with primary antibodies (for instance, anti-cleaved Caspase 3, 1:1000) overnight at 4 °C. After 3 washes with PBS, tissue sections were incubated with biotinylated secondary antibody (1:200 dilution, Vector Laboratories) for 1 h at room temperature then washed thrice, after which streptavidin-horseradish peroxidase conjugates (Vector Laboratories) were added and the slides incubated for 45 min. DAB solution (Vector Laboratories) was then added and the slides were counterstained with haematoxylin.

### Determining senolytic activity

Cells were fixed during days 8-10 after senescence induction and stained with 1 μg/ml DAPI for 15 min to assess cell numbers using automated microscopy. Different models of senescence were used to test senolytic compound activity in cell culture in a similar fashion. For all senescence types induced, a 3-day course of senolytics was applied, with the percentage of cell survival calculated by dividing the number of cells after senolytic treatment against the number of cells treated with vehicle. Alternatively, quantification of cell viability was conducted by collecting all cells and determining the percentage of viable cells by trypan blue exclusion.

### Gene expression analysis by real-time PCR

RNA from cells or tissues was extracted using TRIzol (Life Technologies) and was reverse transcribed to cDNA using the MMLV Reverse Transcriptase kit (Life Technologies) following the manufacturer’s instructions. Real-time PCR was performed using Platinum SYBR Green qPCR SuperMix (Life Technologies) in StepOnePlus Real-Time PCR System (Applied Biosystems). The probes and primers were purchased from Life Technologies. RPL13A was used as an internal control, and primer sequences for relevant genes are listed in Supplementary Table 6.

### Transcriptomic profiling via RNA-seq

Total RNA was prepared from cells using Direct-zol RNA MiniPrep Kit (Zymo Research). RNA sequencing libraries were generated using the NEBNext Ultra RNA Library Prep Kit for Illumina (NEB England BioLabs). Fragmented and randomly primed 2 × 150□bp paired-end libraries were sequenced using Illumina HiSeq X10.

The sequencing data quality were validated by FastQC (version 0.11.8, http://www.bioinformatics.babraham.ac.uk/projects/fastqc/), before aligned to human genome hg38 (or mouse genome mm10) reference genome by TopHat2. For differential gene expression analysis, read count of each gene was obtained by HTSeq (version 0.11.1, https://htseq.readthedocs.io/en/release_0.11.1/). Differentially expressed genes (DEGs) between conditions were identified using DESeq2 R package and genes with Benjamini-Hochberg corrected p-values < 0.05 were defined as differentially expressed. For PCA and unsupervised clustering, the read counts were normalized using rlog from DESeq2 ^64^ and subject to analysis by the Ingenuity Pathways Analysis (IPA) program (http://www.ingenuity.com/index.html). Heatmaps were generated using heatmap.2 function available in gplots R package. Raw data were preliminarily analyzed on the free online platform of Majorbio I-Sanger Cloud Platform (www.i-sanger.com), and subsequently deposited in the NCBI Gene Expression Omnibus (GEO) database under accession codes GSE156301 and GSE164012.

The leading log-fold-change of different conditions calculated by R package edgeR (version 0.38.0, https://bioconductor.org/packages/release/bioc/html/edgeR.html) was used as input for gene set enrichment analysis. The gene sets of mouse molecular signature were obtained by msigdbr (version 7.0.1, https://cran.r-project.org/web/packages/msigdbr/index.html). In this package, the original human genes of Molecular Signatures Database (MSigDB v7.0, http://software.broadinstitute.org/gsea/msigdb/index.jsp) were converted to non-human model organism homologous genes. GSEA was performed using GSEA desktop application version 2.2.7 with Molecular Signature Database (version 3.1), and by clusterProfiler (version 3.14.0) (https://bioconductor.org/packages/release/bioc/html/clusterProfiler.html). R (version 3.6.0, https://cran.r-project.org/) were used for gene expression analysis. The SASP and GSEA signatures were mapped as described before^14^.

Gene ontology (GO) term enrichment analysis was performed using DAVID 6.8 functional annotation tool. The gene lists were selected by comparing gene expression between control and senescent cells or vehicle-treated and drug-treated senescent cells with *t-*test statistics: fold changes >□2 and *P* values□<□0.05. The top GO terms of GO-Biological Progress (BP) and GO-Cellular Component (CC) were selected for presentation.

Protein-protein interaction (PPI) analysis was performed with STRING 3.0. The specific proteins meeting the criteria, were imported to NetworkAnalyst (http://www.networkanalyst.ca). A minimum interaction network was chosen for further hub and module analysis.

### HPLC-qTOF-MS/MS analysis

HPLC-qTOF-MS/MS (high performance liquid chromatograph conjugated with quadrupole time-of-flight-tandem mass spectrometer) analysis was performed on TripleToF 6600 (AB SCIEX) LC-MS/MS. The chromatographic separation was performed on an Accucore C30 column (2.6 um, 250 × 2.1 mm, Thermo) at room temperature. The mobile phase solvent comprises 1% acetic acid in water (solvent A) and 100% methanol (solvent B). The multistep linear gradient solvent system started with 5% B and increasing to 15% B (5 min), 35% B (30 min), 70% B (45 min), 70% B (49 min), 100% B (50 min), held at 100% B for 10 min, and decreasing to 5% B (60 min). At the initial and last gradient step, the column was equilibrated and maintained or washed for 10 min with 5% B. The flow rate was 0.2 ml/min, while the injection volume of samples was 10 ul. Detection was performed using a TripleToF 6600 qTOF mass spectrometer (AB SCIEX) equipped with an ESI interface in negative ion mode. Operation conditions of the ESI source were as follows: capillary voltage, −4000 V; drying gas, 60 (arbitrary units); nebulization gas pressure, 60 psi; capillary temperature, 650 °C; and collision energy, 30. The mass spectra were scanned from 50 to 1200 m/z, with acquisition rate 3 spectra per second. Data acquisition and analysis was performed using the software Analyst TF 1.7.1 (AB SCIEX).

### Radiation of animals

For induction of *in vivo* senescence, C57BL/6J mice (Nanjing Model Animal Centry, China) at 8-12 weeks of age were exposed to a sublethal dose (5 Gy) whole body irradiation (WBI). Eight weeks later, animals were injected with 20 mg/kg PCC1 via intraperitoneal route (*i.p.*) or vehicle once *per* week, for 4 consecutive weeks. Mice were sacrificed 24 h after the last injection. Mouse tissues were harvested for RNA extraction, paraffin embedded for immunohistology, or frozen in OCT solution for cryo-sectioning and SA-β-Gal staining. The mice used for all experiments were randomly assigned to control or treatment groups. Both sexes were used throughout the study.

### Chemotherapeutic regimens

Nod-obese diabetic and severe combined immunodeficiency (NOD-SCID) male mice (Nanjing Model Animal Centry, China) of 6-8 weeks old were housed and maintained in accordance with animal guidelines of Shanghai Institutes for Biological Sciences. For mouse tumor xenograft establishment, human prostate stromal cells (PSC27) and cancer epithelial cells (PC3 or PC3-Luc) were mixed at a ratio of 1:4, with each *in vitro* recombinant comprising 1.25 × 10^6^ total cells prior to subcutaneous implantation as described previously ^14, 37^. Two weeks later, mice were randomized into groups and subject to preclinical treatments. Animals were treated by MIT (0.2 mg/kg) alone, PCC1 (20 mg/kg) along or MIT plus PCC1. Agents were delivered via *i.p.* once *per* biweekly starting from the beginning of the 3rd week, with totally three 2-week cycles throughout the whole regimen as reported formerly ^16, 17^.

Mice were sacrificed at end of an 8^th^ week post tumor xenograft regimen. Primary tumor sizes were measured upon animal dissection, with approximate ellipsoid tumor volume (V) measured and calculated according to the tumor length (l) and width (w) by the formula: V = (π 6) × ((l + w)/2) used previously. Excised tumors were either freshly snap-frozen, or fixed in 10% buffered formalin and subsequently processed as formalin-fixed and paraffin-embedded sections (FFPE) for IHC staining. Tumor growth and metastasis in mice was evaluated using the bioluminescence emitted by PC3-Luc cells which stably express firefly luciferase. The Xenogen IVIS Imager (Caliper Lifesciences) was applied to document BLI across the visible spectrum in isoflurane-anesthetized animals, with the substrate D-Luciferin (150 mg/kg, BioVision) injected subcutaneously each time freshly for imaging-based tumor surveillance.

### Senescent cell clearance and lifespan studies

For aging-related studies, wildtype C57BL/6J male mice were maintained in a specific pathogen-free (SPF) facility at 23-25 °C under a 12 h/12 h light/dark cycle (08:00 to 20:00 light on), with free access to food (standard mouse diet, Lab Diet 5053) and water provided *ad libitum*. The experimental procedure was approved by the Institutional Animal Care and Use Committee (IACUC) at Shanghai Institute of nutritional and health, Chinese Academy of Sciences (SINH, CAS), with all animals handled in accordance with the guidelines for animal experiments defined by IACUC.

Mice were sorted by body weight from low to high, and pairs of mice were selected according to similar body weights. Either control cell (CTRL) or senescent cell (SEN)-transplant, treatments were assigned to every other mouse using a random number generator, with the intervening mice assigned to the other treatment, to allow CTRL- and SEN-transplanted mice matched by weight. After acclimation to the animal facility at SINH, mice aged 5 months were transplanted with cells and treated immediately with vehicle or PCC1 (prepared in ethanol:polyethylene glycol 400:Phosal 50 PG at 10:30:60, 20 mg/kg) for one week (injection via *i.p.* once per 3 days) before subject to bioluminescence imaging (BLI) assays. Physical function tests were performed 1 month after BLI at the age of approximately 25 weeks. The first death occurred approximately 3 months post the last physical function test. For treatment-delayed mice that received cell transplantation, a 1^st^ wave of physical function tests was conducted 5 weeks post cell transplantation. Animals were then subject to treatment with vehicle for 5 days (for those harboring CTRL cells, as the 1^st^ group), vehicle or PCC1 for 5 days (for those harboring SEN cells, as the 2^nd^ and 3^rd^ group, respectively) (each condition via *i.p.* performed consecutively, once per 5 days). Mice were allowed to stay for 2 weeks, after which the 2^nd^ wave of physical function assays was conducted. For senolytics studies associated with natural aging, 20-month-old non-transplanted wildtype C57BL/6J mice were used, which were sorted according to their body weight and randomly assigned to vehicle or PCC1 treatment. Animals were treated once biweekly, in an intermittent manner for 4 months before physical tests were performed at the 24-month age. For senolytics trials pertaining to lifespan extension at advanced age, we used animals at a very old stage. Starting at 24-27 months of age (equivalent to human age of 75-90 years), mice were treated once every 2 weeks (biweekly) with vehicle or PCC1 via oral gavage (20 mg/kg) for 3 consecutive days. Some mice were moved from their original cages during the study to minimize single cage-housing stress. In each case, RotaRod (TSE system, Chesterfield, MO) and hanging tests were chosen for monthly measurement of maximal speed and hanging endurance, respectively, as these tests are considered sensitive and noninvasive. Mice were euthanized and scored as having died if they exhibited more than one of the following signs: (i) unable to drink or eat; (ii) reluctant to move even with stimulus; (iii) rapid weight loss; (iv) severe balance disorder; or (v) bleeding or ulcerated tumor. No mouse was lost due to fighting, accidental death, or dermatitis. The Cox proportional hazard model was used for survival analyses.

### Postmortem pathological examination

Animal cages were checked every day, with dead mice removed from cages. Within 24□h, corpses were opened (abdominal cavity, thoracic cavity, and skull) and preserved in 4% paraformaldehyde (PFA) individually for at least 7 d, with decomposed or disrupted bodies excluded. The preserved bodies were rendered to pathologists for blind examination, with routine assessment procedure followed. Briefly, tumor burden (sum of different types of tumors in each mouse), disease burden (sum of different histopathological changes of major organs in each mouse), severity of each lesion, and inflammation (lymphocytic infiltrate) were evaluated.

### Bioluminescence imaging

Experimental mice were injected via *i.p.* with 3□mg D-luciferin (potassium salt, BioVision Inc) in 200□µl PBS. Mice were anesthetized by inhalation of isofluorane, and bioluminescence images were acquired using a Xenogen IVIS 200 System (Caliper Life Sciences, Hopkinton, MA) according to the manufacturer’s instructions.

### Physical function assessment

All physical tests were performed at least 5 d after the last dose of senolytics treatment. Maximal walking speed was measured using an accelerating RotaRod system (TSE system, Chesterfield, MO). Mice were trained on the RotaRod for 3 d at speeds of 4, 6, and 8 rpm for 200[sec on days 1, 2, and 3. On the test day, mice were placed onto the RotaRod, which was started at 4 rpm. The rotating speed was accelerated from 4 to 40 rpm over a 5-min interval. The speed was recorded when the mouse dropped off the RotaRod, with results averaged from 3 or 4 trials and normalized to baseline speed. Training was not repeated for mice that had been trained within the preceding 2 months. Forelimb grip strength (N) was determined using a Grip Strength Meter (Columbus Instruments), with results averaged over ten trials. To measure grip strength the mouse is swung gently by the tail so that its forelimbs contact the bar. The mouse instinctively grips the bar and is pulled horizontally backwards, exerting a tension. When the tension becomes too great, the mouse releases the bars. For the hanging test, mice were placed onto a 2-mm-thick metal wire that was 35□cm above a padded surface, while animals were allowed to grab the wire with their forelimbs only. Hanging time was normalized to body weight as hanging duration (sec)□×□body weight (g), with results averaged from 2 or 3 trials for each mouse. A Comprehensive Laboratory Animal Monitoring System (CLAMS, Columbus Instruments) was used to monitor daily activity and food intake over a 24-h period (12-h light/12-h dark). The CLAMS system was equipped with an Oxymax Open Circuit Calorimeter System (Columbus Instruments). For treadmill performance, mice were acclimated to a motorized treadmill at an incline of 5° (Columbus Instruments) over 3 d for 5□min each day, starting at a speed of 5□m/min for 2□min and progressing to 7□m/min for 2□min and then 9□m/min for 1□min. On the test day, mice ran on the treadmill at an initial speed of 5□m/min for 2□min, and then the speed was increased by 2□m/min every 2□min until the mice were exhausted. The speed when the mouse dropped from the RotaRod was recorded, and results were averaged from three trials. Exhaustion was defined as the inability to return onto the treadmill despite a mild electrical shock stimulus and mechanical prodding. Distance was recorded and total work (KJ) was calculated using the following formula: mass (kg)□×□g (9.8 m/s^2^) ×□distance (m)□×□sin (5°).

### Comprehensive laboratory animal monitoring system

In a subset of 8-10 mice per group, the habitual ambulatory, rearing, and total activity, oxygen consumption (VO_2_), and carbon dioxide production (VCO_2_) of individual mice were monitored over a 24-h period (12 h light/12 h dark) using a Comprehensive Laboratory Animal Monitoring System (CLAMS) equipped with an Oxymax Open Circuit Calorimeter System (Columbus Instruments). Ambulatory, rearing, and total activity was summed and analyzed for light and dark periods under fed conditions. The VO_2_ and VCO_2_ values were used to calculate the respiratory exchange ratio (RER) and VO_2_. RER values were used to determine the basal metabolic rate (in kilocalories per kilogram *per* hour).

### Tissue SA-**β**-Gal assay and histological evaluation

For SA-β-Gal staining, frozen sections were dried at 37□°C for 20-30□min before fixed for 15□min at room temperature. The frozen sections were washed thrice with PBS and incubated with SA-β-Gal staining solution (Beyotime) overnight at 37□°C. After completion of SA-β-Gal staining, sections were stained with eosin for 1-2□min, rinsed under running water for 1□min, differentiated in 1% acid alcohol for 10-20□s, and washed again under running water for 1□min. Sections were dehydrated in increasing concentrations of alcohol and cleared in xylene. After drying, samples were examined under a bright-field microscope.

Prostate, lung, liver and kidney frozen sections stained with SA-β-Gal were quantified by ImageJ software (NIH) to measure the SA-β-Gal area. The total area was quantified by eosin-positive area, while the relative SA-β-Gal^+^ cells were calculated with the SA-β-G ^+^ area divided by the total area. For the statistics of SA-β-Gal^+^ area of lung, regions of lung were randomly selected to be photographed, avoiding analysis of larger pulmonary blood vessels and trachea. For statistics of SA-β-Gal-positive area of liver, regions were randomly selected to be photographed. For statistics of SA-β-Gal+ area of kidney, regions of renal cortex were randomly selected to be photographed. Each tissue was measured with 10-18 regions.

For endogenous acidic β-galactosidase (β-Gal) staining, the kidney and salivary gland frozen sections were dried at 37□°C for 20-30□min, then fixed in β-Gal staining fix solution for 15□min at room temperature. The frozen sections were washed three times with PBS and incubated with β-Gal staining solution (Beyotime) overnight at 37□°C. The subsequent protocol was similar to that of regular SA-β-Gal staining.

For immunohistochemical (IHC) assays, formalin-fixed paraffin-embedded (FFPE) tissue blocks were cut into 7-µm-thick sections, which were subject to deparaffinization and hydration. Antigen was retrieved in citrate buffer (0.01□M, pH 6.0) at 95□°C for 10□min, and sections were treated with blocking buffer (goat serum 1:60 in 0.1% BSA in PBS) for 60□min at room temperature. Samples were further incubated with Avidin-Biotin (Vector Lab) for 15□min at room temperature. Sections were stained with primary antibody at 4□°C overnight, before incubated with biotinylated secondary antibody (Vector Lab) for 30□min. Finally, fluorescein-labeled avidin DCS (Vector Lab) was applied for 20□min. IHC staining was performed by the pathologist using a BOND RX Fully Automated Research Stainer (Leica). Slides were retrieved for 20□min using Epitope Retrieval 1 (Citrate; Leica) and incubated in Protein Block (Dako) for 5 min. Primary antibodies were diluted in Background Reducing Diluent (Dako) as follows: p16^INK4A^ (rabbit, monoclonal; abcam, Cambridge, MA, catalog no. ab108349) at 1:600, cleaved-caspase 3 (rabbit, polyclonal; Cell Signaling, catalog no. 9661L) at 1:200, and F4/80 (rat, monoclonal; abcam, catalog no. ab90247) at 1:500, except for CD68 antibody (mouse, monoclonal; Dako, catalog no. M0876), which was diluted in Bond Diluent (Leica) at 1:200. All primary antibodies were diluted in Background Reducing Diluent (Dako) and incubated for 15□min, with a Polymer Refine Detection System (Leica) used. This system includes the hydrogen peroxidase block, post primary and polymer reagent, DAB, and hematoxylin. Immunostaining visualization was achieved by incubating the slides for 10□min in DAB and DAB buffer (1:19 mixture) from the Bond Polymer Refine Detection System. Slides were counterstained for 5□min using Schmidt hematoxylin, followed by several rinses in 1□×□Bond wash buffer and distilled water. Slides were subsequently dehydrated with increasing concentrations of ethanol and cleared by xylene, before mounted in xylene-based VECTRASHIELD medium (Vector Lab). Imaging was performed with a LSM 780 Zeiss confocal microscope.

### *In vivo* cytotoxicity assessment via blood test

As routine blood examination, 100 μl fresh blood was acquired from each animal and mixed with EDTA immediately. The blood samples were analyzed by Celltac Alpha MEK-6400 series hematology analyzers (Nihon Kohden). For serum biochemical analysis, blood samples were collected, clotted for 2 h at room temperature or overnight at 4 °C. Samples were then centrifuged (1000 × g, 10 min) to obtain serum. An aliquot of approximately 50 μl serum was subject to analysis for creatinine, urea, alkaline phosphatase (ALP) and alanine transaminase (ALT) by Chemistry Analyzer (Mindray). Evaluation of circulating levels of hemoglobin, white blood cells, lymphocytes and platelets were performed with dry-slide technology on VetTest 8008 chemistry analyzer (IDEXX) as reported previously ^37^.

All animal experiments were conducted in compliance with NIH Guide for the Care and Use of Laboratory Animals (National Academies Press, 2011) and the ARRIVE guidelines, and were approved by the IACUC of Shanghai Institute of Nutrition and Health, Chinese Academy of Sciences (SINH, CAS).

### Measurement of lipid peroxidation

The Levels of 4-hydroxynonenal (HNE)-protein adducts of liver lysates prepared in RIPA buffer were measured in livers of mice using the OxiSelect HNE Adduct Competitive ELISA Kit (Cell Biolabs, San Diego, CA), as formerly described ^65^.

### Examination of glutathione

Mouse livers fixed in 5% sulfosalicylic acid were prepared and analyzed for the concentration of reduced (GSH) and oxidized (GSSG) glutathione using the Glutathione Assay Kit (Cayman Chemical, Ann Arbor, MI), as previously described ^65^. Sample absorbance was measured at wavelength of 405 nm using a plate reader and the ratio of GSH: GSSG was obtained for each tested tissue sample.

### Statistical analysis

Unless otherwise specified, data in the figures are presented as mean ± SD, and statistical significance was determined by unpaired two-tailed Student’s *t*-test (comparing two groups), one-way ANOVA or two-way ANOVA (comparing more than two groups, with Tukey’s post-hoc comparison), Pearson’s correlation coefficients test, Kruskal–Wallis, log-rank test, Wilcoxon–Mann–Whitney test, or Fisher’s exact test with GraphPad Prism 8.3 primed with customized parameters. Cox proportional hazards regression model and multivariate Cox proportional hazards model analysis were performed with statistical software SPSS. Investigators were blinded to allocation during most experiments and outcome assessments. We used baseline body weight to assign mice to experimental groups to achieve similar body weight between groups, so that only randomization within groups matched by body weight was conducted.

To determine sample size, we chose to set the values of type I error (α) and power (1-β) to be statistically adequate: 0.05 and 0.80, respectively ^66^. We then determined N on the basis of the smallest effect we wish to measure. If the required sample size is too large, we chose to reassess the objectives or to more tightly control the experimental conditions to reduce the variance. For all statistical tests, a p value < 0.05 was considered significant, with p values presented as follows throughout the article: ^, not significant; *, *P* < 0.05; **, *P* < 0.01; ***, *P* < 0.001; ****, *P* < 0.0001.

### Data availability

The data that support the findings of this study are available from the corresponding author upon request. The RNA-seq data generated in the present study have been deposited in the Gene Expression Omnibus database under accession numbers GSE156301 and GSE164012.

## Acknowledgements

We are grateful to members of Sun laboratory for reagents, comments and other contributions to this project. The work was supported by grants from National Key Research and Development Program of China (2020YFC2002800, 2016YFC1302400), National Natural Science Foundation of China (NSFC) (81472709, 31671425, 31871380) to Y.S., the Strategic Priority Research Program of Chinese Academy of Sciences (XDB39010500) to Y.S., the fund of Key Laboratory of Tissue Microenvironment and Tumor of Chinese Academy of Sciences (201506, 201706, 202008) to Y.S.; Anti-Aging Collaborative Program of SIBS and BY-HEALTH (C01201911260006) to Y.S., the University and Locality Collaborative Development Program of Yantai (2019XDRHXMRC08) Y.S., and the U.S. DoD PCRP (Idea Development Award PC111703) to Y.S.; U.S. NIH grants R37-AG013925 and P01 AG062413, the Connor Fund, Robert J. and Theresa W. Ryan, and the Noaber Foundation to J.L.K.; National Natural Science Foundation of China (81370730, 81571512) and Yantai Double Hundred Program to Q.F.

## Author contributions

Y.S. conceived this study, designed the experiments and supervised the project. Q.X. performed most of the *in vitro* assays, part of the *in vivo* experiments, and wrote part of the manuscript. Z.L. provided HPLC-qTOF-MS/MS data and conducted preliminary analysis. H.L. performed some cell culture and drug treatment assays. Q.F. performed most of the preclinical studies. X.L, R.H., X.Z., J.C. and J.L.K. provided conceptual inputs or supervised a specific subset of experiments. Y.S. orchestrated data analysis, graphic presentation, and finalized the manuscript. All authors critically read and commented on the final manuscript.

## Competing Interests

The authors declare no competing interests.

## Notes

### Competing Interest Statement

The authors have declared no competing interest.

## References

1. Khosla, S., Farr, J.N., Tchkonia, T. & Kirkland, J.L. The role of cellular senescence in ageing and endocrine disease. Nat Rev Endocrinol 16, 263–275 (2020).

2. Rocca, W.A. et al. Prevalence of multimorbidity in a geographically defined American population: patterns by age, sex, and race/ethnicity. Mayo Clin Proc 89, 1336–1349 (2014).

3. Lopez-Otin, C., Blasco, M.A., Partridge, L., Serrano, M. & Kroemer, G. The hallmarks of aging. Cell 153, 1194–1217 (2013).

4. Pignolo, R.J., Passos, J.F., Khosla, S., Tchkonia, T. & Kirkland, J.L. Reducing Senescent Cell Burden in Aging and Disease. Trends Mol Med 26, 630–638 (2020).

5. Gorgoulis, V. et al. Cellular Senescence: Defining a Path Forward. Cell 179, 813–827 (2019).

6. Baker, D.J. et al. Naturally occurring p16(Ink4a)-positive cells shorten healthy lifespan. Nature 530, 184–189 (2016).

7. Baker, D.J. et al. Clearance of p16Ink4a-positive senescent cells delays ageing-associated disorders. Nature 479, 232–236 (2011).

8. Tchkonia, T. & Kirkland, J.L. Aging, Cell Senescence, and Chronic Disease: Emerging Therapeutic Strategies. JAMA 320, 1319–1320 (2018).

9. Hickson, L.J. et al. Senolytics decrease senescent cells in humans: Preliminary report from a clinical trial of Dasatinib plus Quercetin in individuals with diabetic kidney disease. EBioMedicine 47, 446–456 (2019).

10. Justice, J.N. et al. Senolytics in idiopathic pulmonary fibrosis: Results from a first-in-human, open-label, pilot study. EBioMedicine 40, 554–563 (2019).

11. Zhu, Y. et al. The Achilles’ heel of senescent cells: from transcriptome to senolytic drugs. Aging cell 14, 644–658 (2015).

12. Zhu, Y. et al. Identification of a novel senolytic agent, navitoclax, targeting the Bcl-2 family of anti-apoptotic factors. Aging cell 15, 428–435 (2016).

13. Fuhrmann-Stroissnigg, H. et al. Identification of HSP90 inhibitors as a novel class of senolytics. Nature Communications 8, 422 (2017).

14. Zhang, B.Y. et al. The senescence-associated secretory phenotype is potentiated by feedforward regulatory mechanisms involving Zscan4 and TAK1. Nat Commun 9, 1723 (2018).

15. Sun, Y. et al. Treatment-induced damage to the tumor microenvironment promotes prostate cancer therapy resistance through WNT16B. Nat Med 18, 1359–1368 (2012).

16. Xu, Q. et al. Targeting amphiregulin (AREG) derived from senescent stromal cells diminishes cancer resistance and averts programmed cell death 1 ligand (PD-L1)-mediated immunosuppression. Aging cell 18, e13027 (2019).

17. Han, L. et al. Senescent Stromal Cells Promote Cancer Resistance through SIRT1 Loss-Potentiated Overproduction of Small Extracellular Vesicles. Cancer research 80, 3383–3398 (2020).

18. Chang, J. et al. Clearance of senescent cells by ABT263 rejuvenates aged hematopoietic stem cells in mice. Nat Med 22, 78–83 (2016).

19. Yosef, R. et al. Directed elimination of senescent cells by inhibition of BCL-W and BCL-XL. Nat Commun 7, 11190 (2016).

20. Yousefzadeh, M.J. et al. Fisetin is a senotherapeutic that extends health and lifespan. Ebiomedicine 36, 18–28 (2018).

21. Zhu, Y. et al. New agents that target senescent cells: the flavone, fisetin, and the BCL-XL inhibitors, A1331852 and A1155463. Aging 9, 955–963 (2017).

22. Geng, L., et al. Chemical screen identifies a geroprotective role of quercetin in premature aging. Protein Cell (2018).

23. Hernandez-Segura, A., Nehme, J. & Demaria, M. Hallmarks of Cellular Senescence. Trends in cell biology 28, 436–453 (2018).

24. Chen, B. et al. Curcumin attenuates MSU crystal-induced inflammation by inhibiting the degradation of IkappaBalpha and blocking mitochondrial damage. Arthritis Res Ther 21, 193 (2019).

25. Cadiz-Gurrea, M.D. et al. Cocoa and Grape Seed Byproducts as a Source of Antioxidant and Anti-Inflammatory Proanthocyanidins. International Journal of Molecular Sciences 18 (2017). 26.

26. Rigotti, M. et al. Grape seed proanthocyanidins prevent H2 O2 -induced mitochondrial dysfunction and apoptosis via SIRT 1 activation in embryonic kidney cells. J Food Biochem 44, e13147 (2020).

27. Long, M. et al. The Protective Effect of Grape-Seed Proanthocyanidin Extract on Oxidative Damage Induced by Zearalenone in Kunming Mice Liver. Int J Mol Sci 17 (2016).

28. Chen, F. et al. Grape seed proanthocyanidin inhibits monocrotaline-induced pulmonary arterial hypertension via attenuating inflammation: in vivo and in vitro studies. J Nutr Biochem 67, 72–77 (2019).

29. Georgieva, E. et al. Mitochondrial Dysfunction and Redox Imbalance as a Diagnostic Marker of “Free Radical Diseases”. Anticancer research 37, 5373–5381 (2017).

30. Zhu, W., Li, M.C., Wang, F.R., Mackenzie, G.G. & Oteiza, P.I. The inhibitory effect of ECG and EGCG dimeric procyanidins on colorectal cancer cells growth is associated with their actions at lipid rafts and the inhibition of the epidermal growth factor receptor signaling. Biochem Pharmacol 175, 113923 (2020).

31. Koteswari, L.L., Kumari, S., Kumar, A.B. & Malla, R.R. A comparative anticancer study on procyanidin C1 against receptor positive and receptor negative breast cancer. Nat Prod Res 34, 3267–3274 (2020).

32. van der Feen, D.E. et al. Cellular senescence impairs the reversibility of pulmonary arterial hypertension. Sci Transl Med 12 (2020).

33. Aguayo-Mazzucato, C. et al. Acceleration of beta Cell Aging Determines Diabetes and Senolysis Improves Disease Outcomes. Cell Metab 30, 129–142 e124 (2019).

34. Khan, S. et al. A selective BCL-XL PROTAC degrader achieves safe and potent antitumor activity. Nat Med 25, 1938–1947 (2019).

35. Sun, P. et al. Trimer procyanidin oligomers contribute to the protective effects of cinnamon extracts on pancreatic beta-cells in vitro. Acta Pharmacol Sin 37, 1083–1090 (2016).

36. Gao, W. et al. Procyanidin B1 promotes in vitro maturation of pig oocytes by reducing oxidative stress. Mol Reprod Dev 88, 55–66 (2021).

37. Chen, F. et al. Targeting SPINK1 in the damaged tumour microenvironment alleviates therapeutic resistance. Nat Commun 9, 4315 (2018).

38. Melisi, D. et al. Modulation of pancreatic cancer chemoresistance by inhibition of TAK1. Journal of the National Cancer Institute 103, 1190–1204 (2011).

39. Xu, M. et al. Senolytics improve physical function and increase lifespan in old age. Nat Med 24, 1246–1256 (2018).

40. Stoupi, S. et al. In vivo bioavailability, absorption, excretion, and pharmacokinetics of [14C]procyanidin B2 in male rats. Drug metabolism and disposition: the biological fate of chemicals 38, 287–291 (2010).

41. Jorgensen, E.M., Marin, A.B. & Kennedy, J.A. Analysis of the oxidative degradation of proanthocyanidins under basic conditions. J Agric Food Chem 52, 2292–2296 (2004).

42. Partridge, L., Fuentealba, M. & Kennedy, B.K. The quest to slow ageing through drug discovery. Nat Rev Drug Discov (2020).

43. Liu, Y. et al. Expression of p16(INK4a) in peripheral blood T-cells is a biomarker of human aging. Aging cell 8, 439–448 (2009).

44. Zeng, Y. et al. Comparison of In Vitro and In Vivo Antioxidant Activities of Six Flavonoids with Similar Structures. Antioxidants (Basel) 9 (2020).

45. Rodriguez-Ramiro, I., Martin, M.A., Ramos, S., Bravo, L. & Goya, L. Comparative effects of dietary flavanols on antioxidant defences and their response to oxidant-induced stress on Caco2 cells. Eur J Nutr 50, 313–322 (2011).

46. He, S. & Sharpless, N.E. Senescence in Health and Disease. Cell 169, 1000–1011 (2017).

47. Amor, C. et al. Senolytic CAR T cells reverse senescence-associated pathologies. Nature 583, 127–132 (2020).

48. Childs, B.G. et al. Senescent intimal foam cells are deleterious at all stages of atherosclerosis. Science 354, 472–477 (2016).

49. Jeon, O.H. et al. Local clearance of senescent cells attenuates the development of post-traumatic osteoarthritis and creates a pro-regenerative environment. Nat Med 23, 775–781 (2017).

50. Riessland, M. et al. Loss of SATB1 Induces p21-Dependent Cellular Senescence in Post-mitotic Dopaminergic Neurons. Cell Stem Cell 25, 514–530 e518 (2019).

51. Hou, Y. et al. Ageing as a risk factor for neurodegenerative disease. Nat Rev Neurol 15, 565–581 (2019).

52. Wang, Y.Y. et al. Discovery of piperlongumine as a potential novel lead for the development of senolytic agents. Aging-Us 8, 2915–2926 (2016).

53. Guerrero, A. et al. Cardiac glycosides are broad-spectrum senolytics. Nat Metab 1, 1074–1088 (2019).

54. Triana-Martinez, F. et al. Identification and characterization of Cardiac Glycosides as senolytic compounds. Nature Communications 10, 4731 (2019).

55. Childs, B.G. et al. Senescent cells: an emerging target for diseases of ageing. Nat Rev Drug Discov 16, 718–735 (2017).

56. Schafer, M.J. et al. Cellular senescence mediates fibrotic pulmonary disease. Nat Commun 8, 14532 (2017).

57. Cai, Y. et al. Elimination of senescent cells by beta-galactosidase-targeted prodrug attenuates inflammation and restores physical function in aged mice. Cell research, 1–16 (2020).

58. Zhu, Y. et al. New agents that target senescent cells: the flavone, fisetin, and the BCL-X-L inhibitors, A1331852 and A1155463. Aging-Us 9, 955–963 (2017).

59. Zhao, J., Wang, J., Chen, Y. & Agarwal, R. Anti-tumor-promoting activity of a polyphenolic fraction isolated from grape seeds in the mouse skin two-stage initiation-promotion protocol and identification of procyanidin B5-3’-gallate as the most effective antioxidant constituent. Carcinogenesis 20, 1737–1745 (1999).

60. Nie, Y. & Sturzenbaum, S.R. Proanthocyanidins of Natural Origin: Molecular Mechanisms and Implications for Lipid Disorder and Aging-Associated Diseases. Adv Nutr 10, 464–478 (2019).

61. Bae, J., Kumazoe, M., Murata, K., Fujimura, Y. & Tachibana, H. Procyanidin C1 Inhibits Melanoma Cell Growth by Activating 67-kDa Laminin Receptor Signaling. Molecular nutrition & food research 64, e1900986 (2020).

62. Ma, H. et al. Macrophages inhibit adipogenic differentiation of adipose tissue derived mesenchymal stem/stromal cells by producing pro-inflammatory cytokines. Cell Biosci 10, 88 (2020).

63. Finney, D.J. The adjustment for a natural response rate in probit analysis. Ann Appl Biol 36, 187–195 (1949).

64. Love, M.I., Huber, W. & Anders, S. Moderated estimation of fold change and dispersion for RNA-seq data with DESeq2. Genome biology 15, 550 (2014).

65. Robinson, A.R. et al. Spontaneous DNA damage to the nuclear genome promotes senescence, redox imbalance and aging. Redox Biol 17, 259–273 (2018).

66. Krzywinski, M. & Altman, N. POINTS OF SIGNIFICANCE Power and sample size. Nat Methods 10, 1139–1140 (2013).

