## Supplemental Figures and Legends for "Procyanidin C1 is a natural agent with senolytic activity against aging and age-related diseases"

### Extended Data Fig. 1

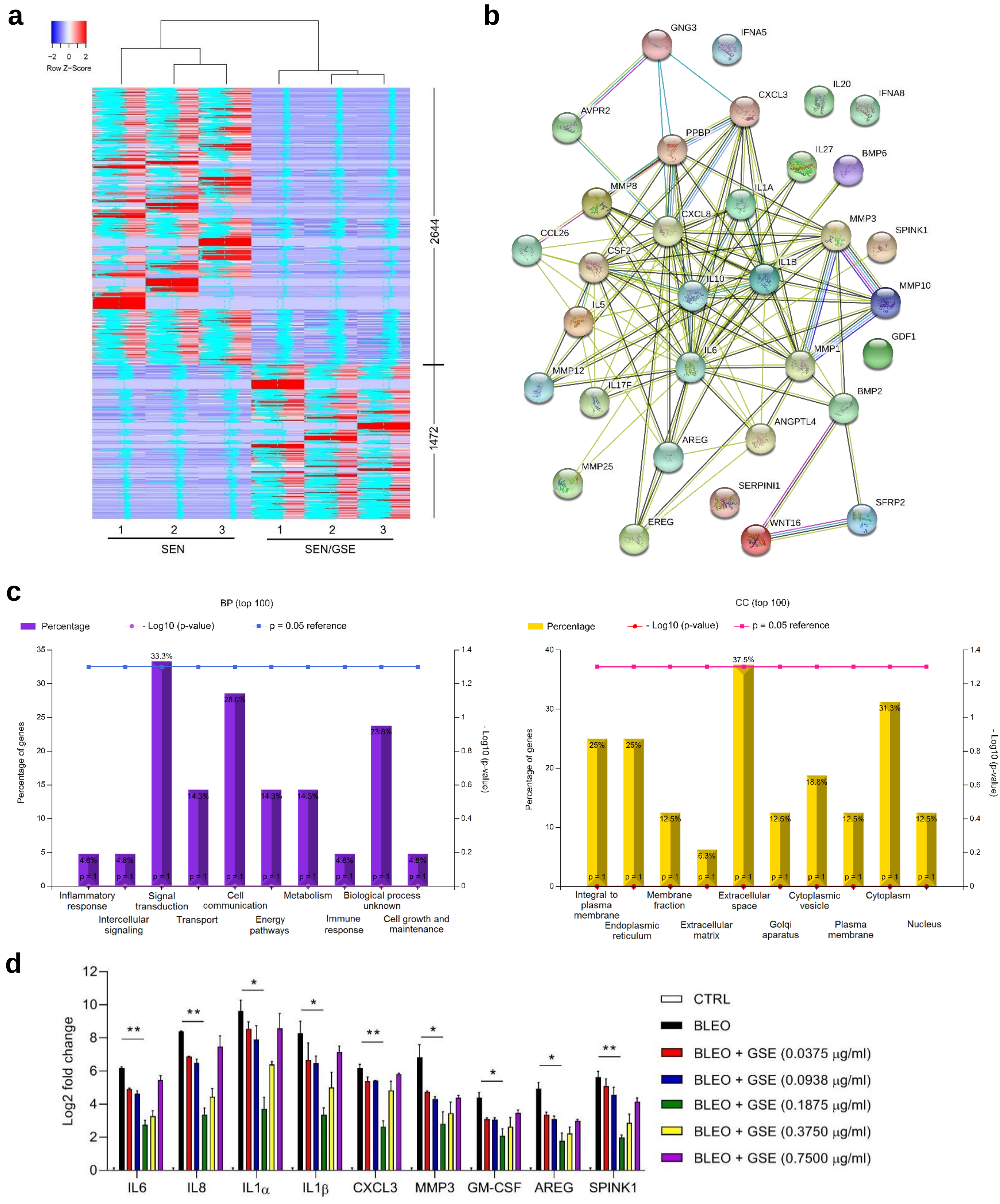

**Extended Data Fig. 2**

**a**

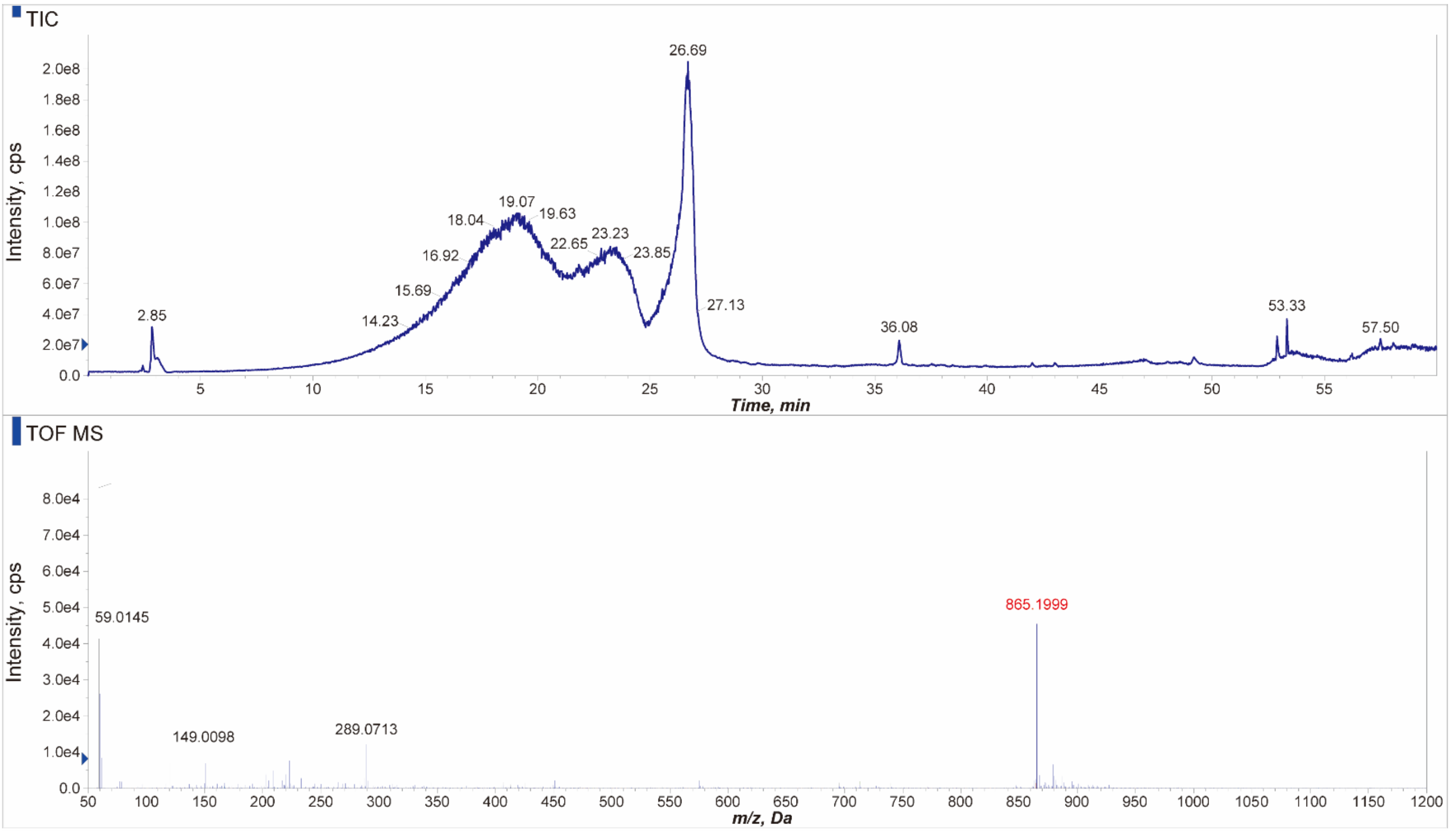

### Extended Data Fig. 3

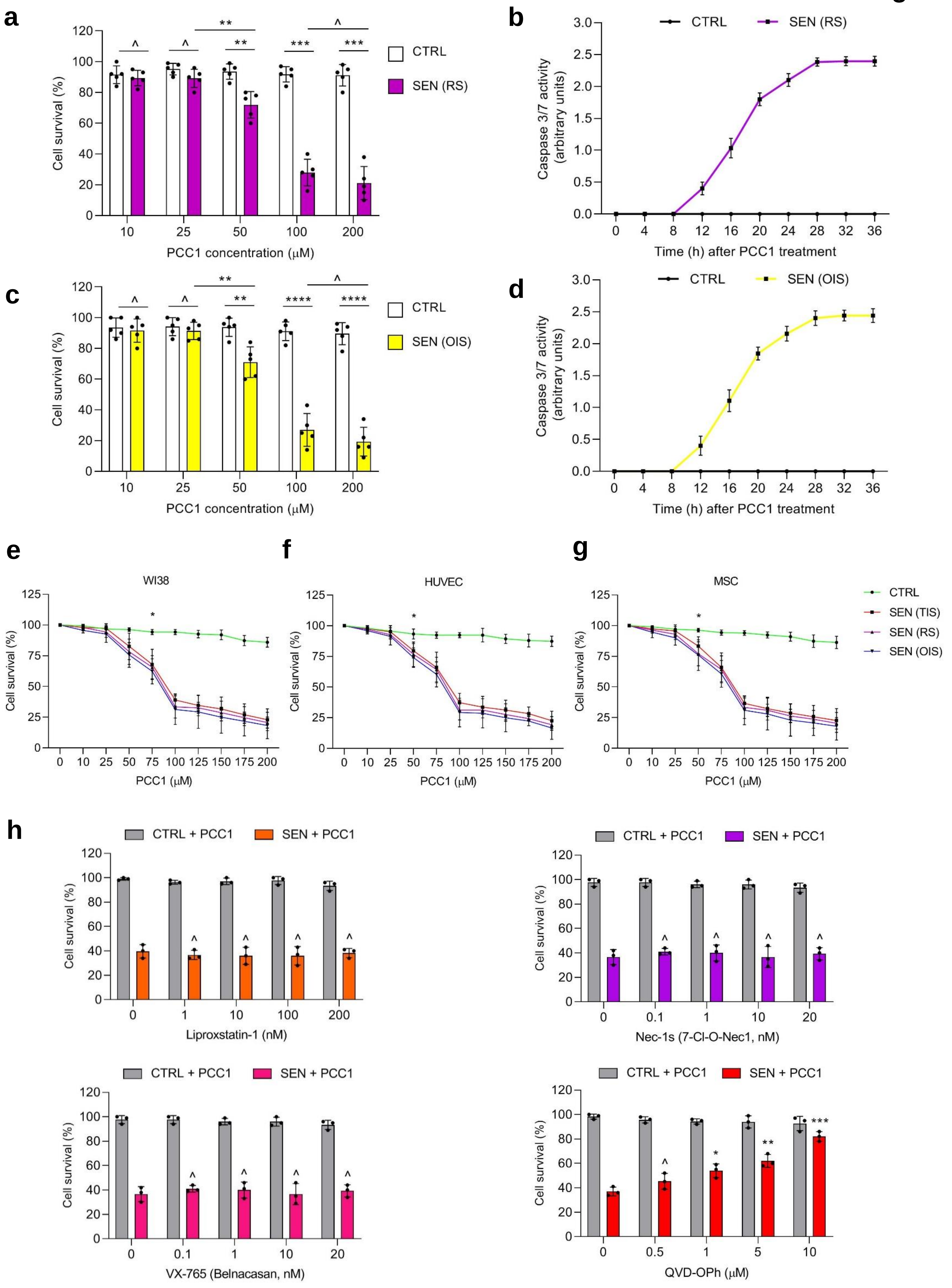

### Extended Data Fig. 4

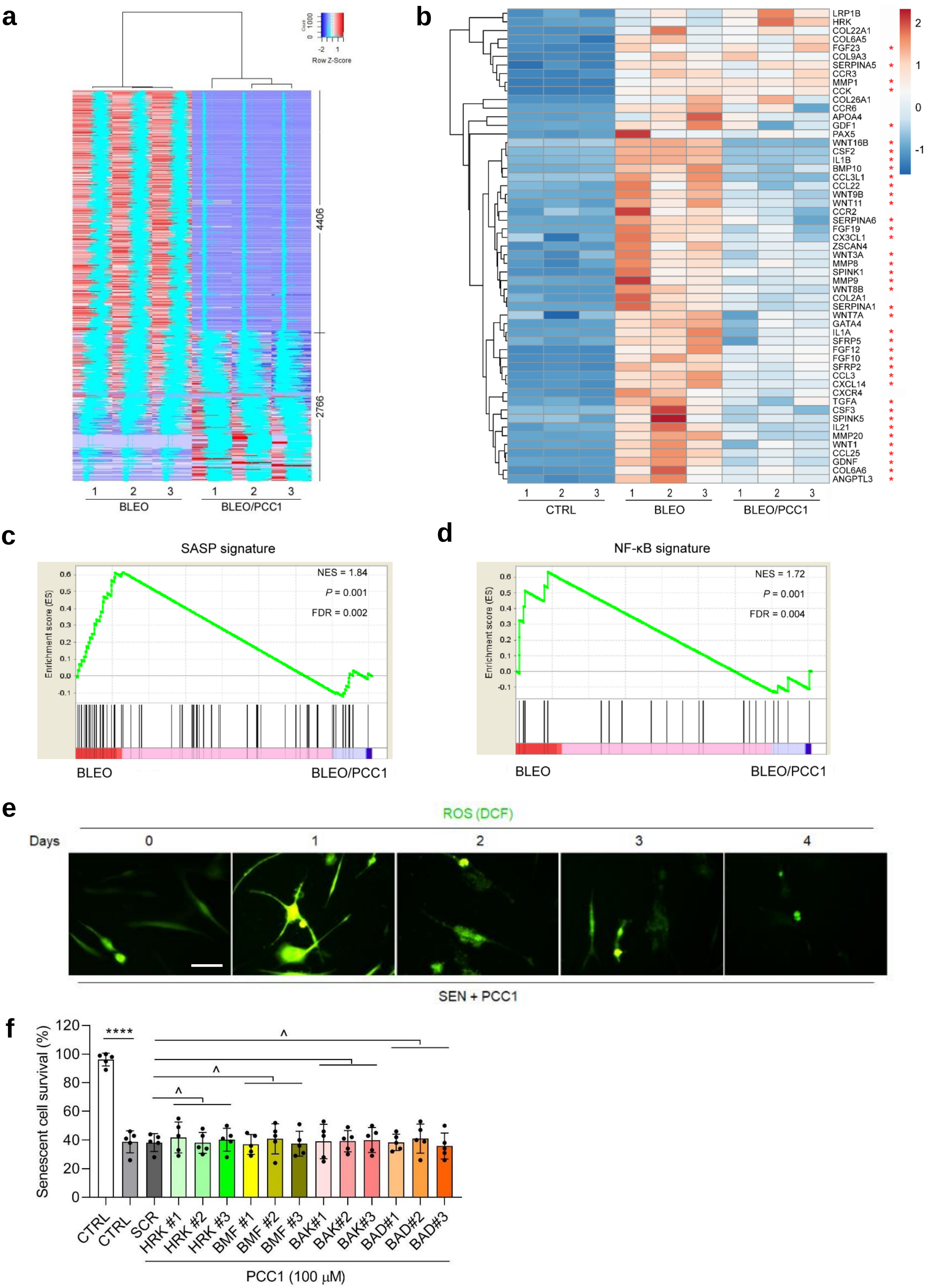

### Extended Data Fig. 5

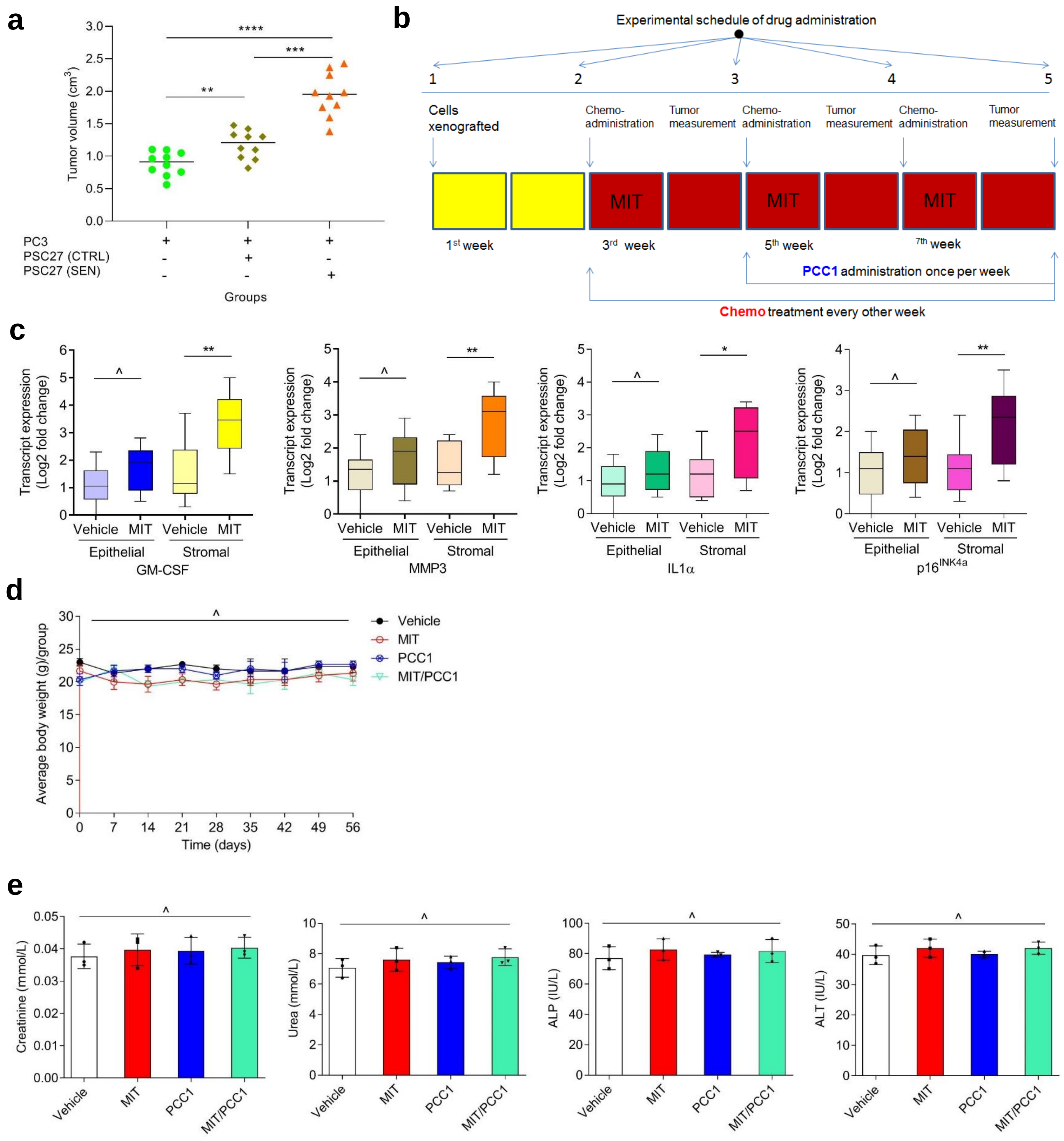

### Extended Data Fig. 6

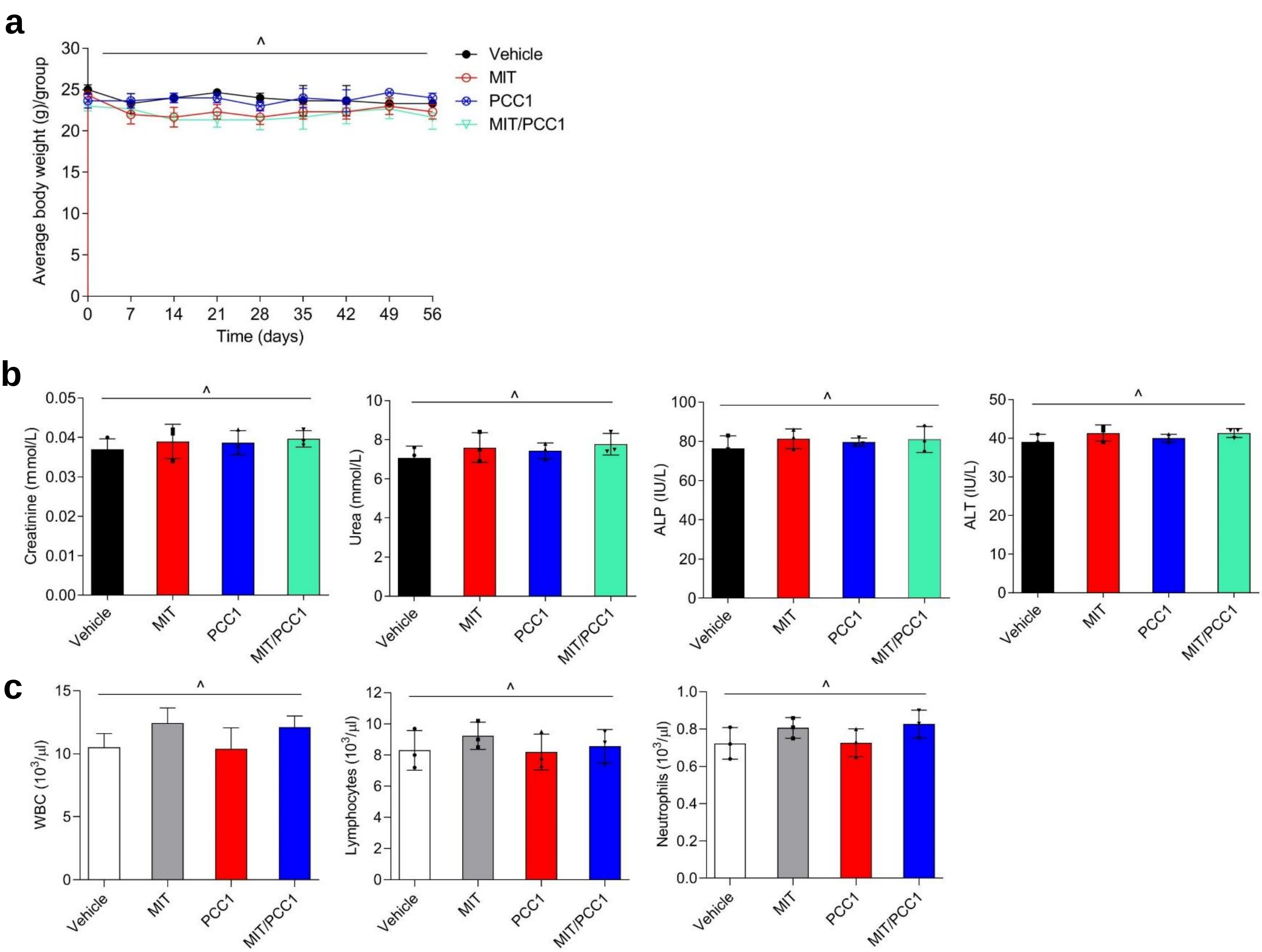

### Extended Data Fig. 7

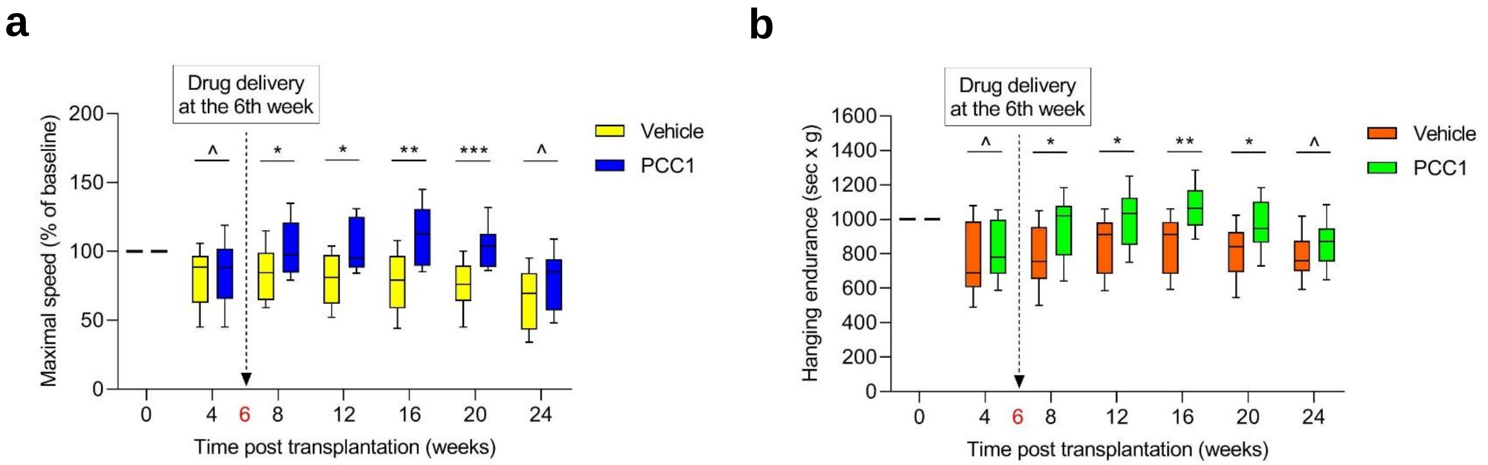

### Extended Data Fig. 8

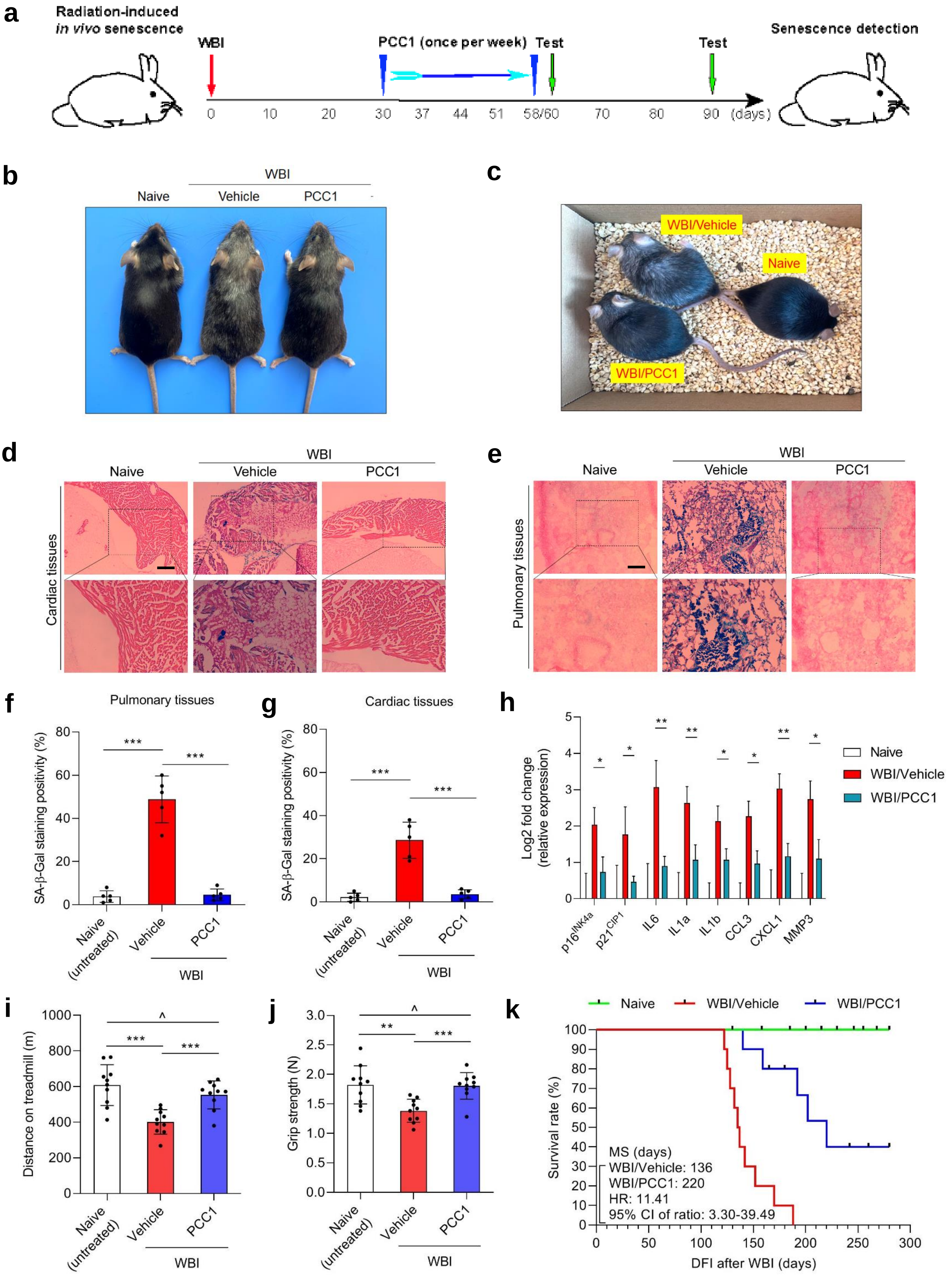

Extended Data Fig. 9

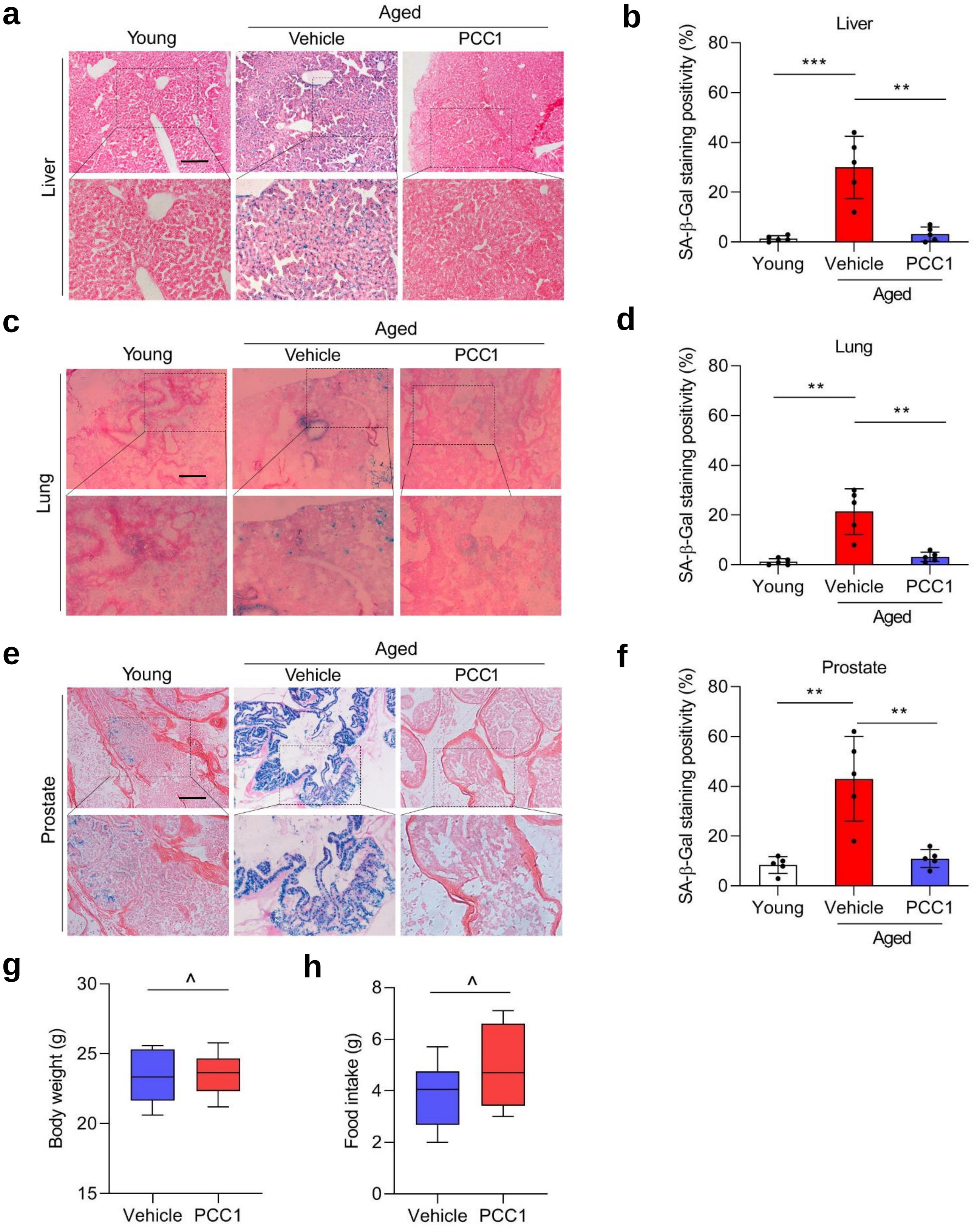

**Extended Data Fig. 10**

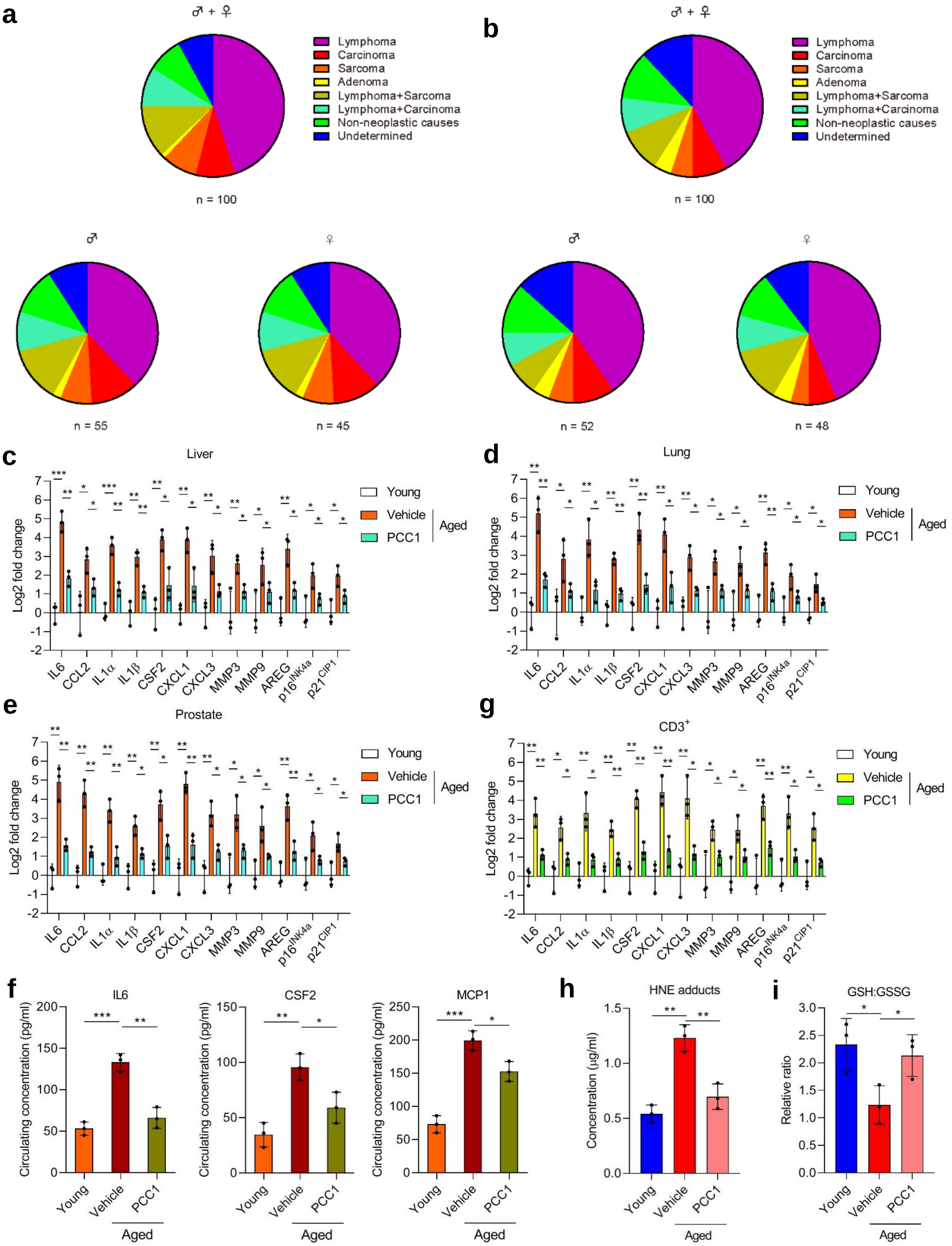

#### Extended Data Figure Legends

**Extended Data Fig. 1. Bioinformatics profiling of the effect of GSE on senescent cells.**

(**a**) Heatmap depicting the expression landscape of senescent cells and their counterparts exposed to GSE. Note there were 2644 and 1472 genes, expression of which was significantly downregulated and upregulated by GSE, respectively. (**b**) Mapping of protein-protein interaction (PPI) for BLEO-upregulated but GSE-downregulated genes by STRING program. (**c**) Column charts depicting the biological processes (BP, left) and cellular components (CC, right) pronouncedly associated with transcripts upregulated by BLEO but downregulated by GSE as revealed by GO analysis. (**d**) Quantitative analysis of SASP factor expression at transcription level when BLEO-induced senescent cells were subject to treatment by different concentrations of GSE in culture. *, $P < 0.05$; **, $P < 0.01$. Data in **d** are representative of 3 independent biological replicates.

**Extended Data Fig. 2. HPLC Mass spectrum profiling of PCC1 by analytical HPLC.**

(**a**) High resolution mass spectra showing the total ion chromatogram (TIC, upper) and base peak chromatogram (BPC, lower) of procyanidin C1 (PCC1) after performance of HPLC-ESI-qTOF-MS.

**Extended Data Fig. 3. *In vitro* cell assays identify PCC1 as a broad-spectrum senolytic.**

(**a**) Senolytic activity appraisal by measuring the percentage of surviving senescent PSC27 cells induced by replicative exhaustion (RS) at increasing PCC1 concentrations. (**b**) Caspase 3/7 activity-based apoptotic evaluation of CTRL and RS cells treated by PCC1. (**c**) Senolytic activity assessment by determining the percentage of surviving senescent cells induced by oncogenic $HRas^{G12V}$ (OIS) at increasing PCC1 concentrations. (**d**) Caspase 3/7 activity-based apoptotic appraisal of CTRL and OIS cells treated by PCC1. (**e-g**) Quantification of the viability of cells in CTRL, TIS, RS and OIS groups treated by 100 μM for 3 d. Left, WI38. Middle, HUVEC. Right, MSC. (**h**) Quantification of cell survival in CTRL and SEN populations after combined treatment of 100 μM PCC1 with a ferroptosis inhibitor (liproxstatin-1), necroptosis inhibitor (Nec-1s), caspase-1 inhibitor (VX-765), or a pan-caspase inhibitor (QVD-O-Ph). Cell survival was presented as comparative data of different concentrations of chemicals relative to cells treated by the vehicle. All error bars represent mean ± SD, and data are representative of 3 biological replicates. ^, $P > 0.05$. *, $P < 0.05$. **, $P < 0.01$. ***, $P < 0.001$. ****, $P < 0.0001$.

**Extended Data Fig. 4. Bioinformatics profiling of the effect of PCC1 on expression of senescent cells.**

(**a**) Heatmap displaying human genes, the expression of which was significantly upregulated (4406) or downregulated (2766) by PCC1 in senescent PSC27 cells. (**b**) Heatmap depicting top genes (50) which were significantly upregulated in senescent cells but downregulated upon treatment by PCC1. (**c**) GSEA blot of a significant gene set of the SASP spectrum. (**d**) GSEA blot of significant gene set associated with signaling mediated by the NF-κB pathway. (**e**) Staining of reactive oxygen species (ROS) in senescent PSC27 cells with 2'-7'-dichlorodihydrofluorescein diacetate (DCFH-DA), a cell permeable fluorescent probe indicative of changes in the redox state. Signals were measured in a 3-d time course starting from PCC treatment. (**f**) Survival assays of PSC27 cells that were infected with lentivirus encoding shRNAs targeting HRK, BMF, BAK or BAD, individually, prior to senescence induction. Cells were then exposed to PCC1 (100 μM) 7 days after BLEO-mediate damage, for a 3-d period to induce apoptosis. SCR, scramble shRNA. Data of bar graphs are shown as mean ± SD and representative of 3 biological replicates. ^, $P > 0.05$. ****, $P < 0.0001$.

**Extended Data Fig. 5. Schematic design of preclinical trial, expression analysis of the SASP and pathophysiological appraisal of treatment effects.**

(**a**) Statistical analysis of tumor end volumes. PC3 cells were xenografted alone or together with PSC27 cells to the hind flank of animals, with tumor volumes measured at the end of an 8-week period. (**b**) Cancer cells (PC3) alone or alongside stromal cells (PSC27) were inoculated subcutaneously to NOC/SCID mice 2 weeks prior to chemotherapy. MIT was provided via intravenous injection on the 1$^{st}$ day of each week starting from the 3$^{rd}$ week, then given every other week with a total number of 3 doses. PCC1 was delivered via i.p. starting from the beginning of the 5$^{th}$ week, then on an every-other-week schedule. At the end of 8 weeks mice were sacrificed, tumor volume measured and tissues histologically assessed. (**c**)Transcript analysis of several canonical SASP factors including GM-CSF, MMP3 and IL1α, and p16$^{INK4a}$ expressed in the epithelial and stromal cells, respectively. Individual cell types were isolated from tumor tissues via LCM. (**d**) Body weight determination performed on a weekly basis for immunodeficient mice. (**e**) Serum measurement of creatinine, urea, alkaline phosphatase (ALP), and alanine aminotransferase (ALT) with terminal bleeds (cardiac punctures) taken at the end of therapeutic regimens. Data are shown as mean ± SD and representative of 3 independent experiments. MIT, mitoxantrone. N = 10 per treatment arm. ^, $P > 0.05$. *, $P < 0.05$. **, $P < 0.01$. ***, $P < 0.001$. ****, $P < 0.0001$.

**Extended Data Fig. 6. Pathophysiological assessment of treatment effects on immunocompetent mice.**

(**a**) Animal body weight determination performed on a weekly basis for immunocompetent C57BL/6J mice. (**b**) Serum measurement of creatinine, urea, alkaline phosphatase (ALP), and alanine aminotransferase (ALT) with terminal bleeds (cardiac punctures) taken at the end of therapeutic regimens. (**c**) Routine analysis of peripheral blood. The circulating levels of hemoglobin, white blood cells, lymphocytes and platelets at the end of each therapeutic regimen were assessed. Data are shown as mean ± SD and representative of 3 independent experiments. MIT, mitoxantrone. WBC, white blood count. N = 3 per treatment arm. ^, $P > 0.05$.

**Extended Data Fig. 7. PCC1 alleviates physical dysfunction of animals transplanted with senescent cells.**

(**a**) Maximal walking speed (relative to the baseline) of 5-month-old male C57BL/6J mice at different time points post transplantation of $1 \times 10^6$ SEN stromal cells and treatment with PCC1 or vehicle (V). (**b**) Hanging endurance (measured as sec x g) of 5-month-old male C57BL/6J mice at different time points as described in (**a**). Data are shown as mean ± SD and representative of 3 independent experiments. ^, $P > 0.05$. *, $P < 0.05$. **, $P < 0.01$. ***, $P < 0.001$.

**Extended Data Fig. 8. Clearing senescent cells by PCC1 alleviates physical dysfunction of animals exposed to WBI.**

(**a**) Schematic presentation of experimental procedure for mouse experiencing whole body irradiation (WBI) and physical function measurements. (**b**) Whole body snapshoot comparison of C57BL/6J mice that were naïve, WBI-exposed followed by vehicle-treatment and WBI-exposed but PCC1-treated, respectively. (**c**) An in-cage picture for animals described in (**a**) preclinical conditions. (**d**) Representative images of SA-β-Gal staining of cardiac tissues of untreated (naive) and WBI-treated mice subject to vehicle or PCC1 treatment. Scale bar,

200 μm. (**e**) Representative images of SA-β-Gal staining of pulmonary tissues of mice as described in (**d**). Scale bar, 200 μm. (**f**) Comparative statistics of SA-β-Gal staining positivity of cardiac tissues of animals examined in (**d**). (**g**) Comparative statistics of SA-β-Gal staining positivity of pulmonary tissues of animals examined in (**e**). (**h**) Quantitative measurement of SASP factor expression at transcription level in tissues collected from animals treated in conditions of (**a**). (**i-j**) Measurement of running distance on treadmill (**i**) and grip strength (**j**) for experimental mice. (**k**) Kaplan-meier survival analysis of C57BL/6J mice exposed to WBI and treated weekly with vehicle or PCC1, with naïve examined as untreated control. Data are shown as mean ± SD and representative of 3 independent experiments. ^, $P > 0.05$. *, $P < 0.05$. **, $P < 0.01$. ***, $P < 0.001$. ****, $P < 0.0001$.

**Extended Data Fig. 9. Intermittent administration of PCC1 alleviates physical dysfunction of aged mice.**

(**a**) Representative images of SA-β-Gal staining of liver tissues from young (6 months of age) and aged mice treated with vehicle or PCC1. Scale bar, 200 μm. (**b**) Comparative statistics of SA-β-Gal staining positivity of samples assayed in (**a**). (**c**) Representative images of SA-β-Gal staining of lung tissues from young and aged mice treated with vehicle or PCC1. Scale bar, 200 μm. (**d**) Comparative statistics of SA-β-Gal staining positivity of samples examined in (**a**). (**e**) Representative images of SA-β-Gal staining of prostate tissues from young and aged mice treated with vehicle or PCC1. Scale bar, 200 μm. (**f**) Comparative statistics of SA-β-Gal staining positivity of samples examined in (**e**). (**g**) Quantification of body weight of animals as described in (**a**). (**h**) Measurement of food intake of animals as described in (**a**). Data are shown as mean ± SD and representative of 3 independent experiments. ^, $P > 0.05$. *, $P < 0.05$. **, $P < 0.01$. ***, $P < 0.001$.

**Extended Data Fig. 10. Late life intervention with PCC1 does not alter the**

**cause of death, but restrains the SASP and reduces oxidative stress.**

(**a**-**b**) Pie charts depicting the ultimate causes of death of C57BL/6J mice that had undergone vehicle (**a**) or PCC1 (**b**) biweekly treatment starting from 24-27 months of age. Note there was no significant difference between vehicle- and PCC1-treated groups upon analysis using either Chi-square or Fisher’s exact tests. (**c**-**e**) qRT-PCR profiling of SASP and senescence marker expression in tissues of solid organs, including liver (**c**), lung (**d**) and prostate (**e**) collected from young (6-month-old, untreated), aged (24-27-month-old) vehicle-treated and aged (24-27-month-old) PCC1-treated animals, respectively. (**f**) Measurement of circulating levels of hallmark SASP factors IL6, CSF2 and MCP-1 in mouse blood by ELISA assays. (**g**) Quantification of SASP and senescence marker expression in $CD3^+$ peripheral T cells of experimental mice described in **c**-**f**. (**h**) Examination of 4-hydroxynonenal (HNE) adducts, a marker of lipid peroxidation and oxidative stress by ELISA measurement with tissue lysates of the liver. (**i**) Determination of the ratio of reduced (GSH) to oxidized (GSSG) glutathione measured as an index oxidative stress. N = 3 *per* group for assays of **c**-**h**. Data represented as mean ± SD and are representative of 3 independent biological replicates. Unpaired two-tailed Student's *t*-test. ^, $P > 0.05$. *, $P < 0.05$. **, $P < 0.01$. ***, $P < 0.001$.
